# Combination therapy targeting inflammasome and fibrogenesis alleviates inflammation and fibrosis in a zebrafish model of silicosis

**DOI:** 10.1101/2022.05.04.490486

**Authors:** Sylwia D. Tyrkalska, Annamaria Pedoto, Alicia Martínez-López, Sergio Candel, José A. Ros-Lucas, Pablo Mesa-del-Castillo, Victoriano Mulero

**Affiliations:** Departmento de Biología Celular e Histología, Facultad de Biología, Universidad de Murcia, 30100 Murcia, Spain; Instituto Murciano de Investigación Biosanitaria (IMIB)-Arrixaca, 30120 Murcia, Spain; Centro de Investigación Biomédica en Red de Enfermedades Raras (CIBERER), Instituto de Salud Carlos III, 28029 Madrid, Spain; Servicio de Neumología, Hospital Clínico Universitario Virgen de la Arrixaca, 30120 Murcia, Spain; Servicio de Reumatología, Hospital Clínico Universitario Virgen de la Arrixaca, 30120 Murcia, Spain

**Keywords:** silicosis, silica crystals, fibrosis, inflammation, TLRs, inflammasome, zebrafish

## Abstract

Silicosis is a long-term lung disease caused by the inhalation of large amounts of crystalline silica dust. As there is no effective treatment available, patients are provided with supportive care, and some may be considered for lung transplantation. There is therefore an evident need for a better understanding of the disease’s biology and for identifying new therapeutic targets and therapies. In this context, our group has developed a larval zebrafish model of silicosis by injecting silica crystals into the hindbrain ventricle, a cavity into which immune cells can be recruited and that mimics the alveolar environment of the human lung. The injection of silica crystals into this cavity led to the initiation of local and systemic immune responses driven through both TLR- and inflammasome-dependent signaling pathways, followed by fibrosis, as happens in human patients. The combination of the inflammasome inhibitor VX-765 and the antifibrotic agent pirfenidone was found to be the best therapy to alleviate both inflammation and fibrosis. The zebrafish model of silicosis developed here is a unique tool that will shed light onto the molecular mechanisms involved in the progression of this devastating disease and for identifying novel drugs that improve the quality of life of silicosis patients.

## INTRODUCTION

Silicosis, is a fibrotic lung disease caused by the inhalation of free crystalline silicon dioxide or silica and which may be regarded as a preventable occupational respiratory disease (Leung et al., 2012). The disease is classified as pneumoconiosis, an interstitial lung disease caused by the inhalation of dust, usually in a work environment, causing interstitial fibrosis and scarring inside the lung. Silicosis is marked by inflammation and fibrotic nodular lesions in the upper lobes of the lungs. It is progressive disease and is characterized by shortness of breath, cough, chest pain, wheezing, fever, and cyanosis, which is why it may often be misdiagnosed as pulmonary edema, pneumonia, or even tuberculosis (Rees and Murray, 2007). The most common form of the disease is called chronic silicosis, which develops 10 or more years after exposure to relatively low levels of respirable crystalline silica, even after occupational exposure has stopped (Rushton, 2007). Sometimes, symptoms may appear within 5-10 years of exposure, especially in people who have been exposed to large amounts of silica dust, in which case it is called accelerated silicosis (Barnes et al., 2019; Edwards, 2019). Finally, very high levels of exposure might cause symptoms within weeks or months, causing the most severe type of the disease, known as acute silicosis (Barnes et al., 2019; Duchange et al., 1998).

Silicon (Si) is the second most common element in the earth’s crust, after oxygen. Both of those elements are able to react with each other and form the compound called silica, also known as silicon dioxide (SiO_2_) (Martin, 2007). As oxygen and silicon make up almost 75% of the earth’s crust, the presence of silica is very common (Kramer et al., 2012). It can be found in many rocks, metals, soil and even forms part of sand. Silica may appear in three different forms depending on the structure of the molecule, the one responsible for silicosis is the crystalline form (less than 10 µm in diameter), which depending on the temperature of formation, can exist in seven different sub-forms, the most well known being quartz, cristobalite, and tridymite (Carrieri et al., 2020). Respirable crystalline silica may be generated during mechanical operations like cutting, grinding, polishing, or crushing of stone, rock, concrete, brick, and mortar (Barnes et al., 2019).

Looking deeper into the molecular basis of silicosis, the underlying cause of the disease is the inhalation of respirable silica particles, which can easily reach the lower respiratory tract and the oxygen and carbon dioxide exchange zones. From here, the lungs are unable to clear out the dust via mucous or coughing. The silica particles can embed themselves deeply in the tiny alveolar sacs and ducts, where they are recognized by receptors localized on the surface of alveolar macrophages and, after being phagocytized by them, they may persist at the site and trigger inflammatory process (Lopes-Pacheco et al., 2016; Mossman and Churg, 1998). The receptor most associated with silica binding are scavenger receptors (SR), the transmembrane proteins SR-AI and SR-AII, macrophage receptors with a collagenous structure (MARCO), toll-like receptor 4 (TLR4), and type 3 complement receptor (Chan et al., 2018; Hamilton et al., 2008; Palecanda and Kobzik, 2001). It is worth noting that silica is extremely toxic for macrophages; however, if macrophages survive contact with silica particles, they can migrate out of the lung or move to the lung interstitium, where they convert into activated interstitial macrophages (Bowden et al., 1989; Lapp and Castranova, 1993). The internalized silica particles get entrapped by lysosomes, in which a variety of enzymes are prepared to digest the particles. However, the silica particle cannot be broken down by the enzymes, which results in the loss of lysosomal membrane integrity and the release of lysosomal enzymes, such as cathepsin B, which activates cell apoptosis and the NLRP3 inflammasome (Riteau et al., 2012; Thibodeau et al., 2004) (Cassel et al., 2008; Harijith et al., 2014; Peeters et al., 2013; Sayan and Mossman, 2016). Moreover, the inflammatory response provokes the release of tumor necrosis factors (TNFs), interleukin-1 (IL-1), leukotriene B4 (LTB4) and other cytokines as well as the production of reactive oxygen species (ROS) (Mossman and Churg, 1998). In turn, these insults stimulate fibroblasts for proliferation and the production of collagen around the silica particles. All this damages the pulmonary parenchyma, and the subsequent repair and regeneration processes may result in fibrosis and the formation of nodular lesions or even lead to carcinogenesis (Leung et al., 2012; Lopes-Pacheco et al., 2016; Mossman and Churg, 1998).

Although, some animal models are already available to study silicosis, namely mice and rats, there is still a need to develop new *in vivo* animal models to study the pathogenesis and the molecular basis of this disease, and for developing drugs and treatments (Cao et al., 2020; Honnons and Porcher, 2000; Mayeux et al., 2019). During the last 20 years the zebrafish has emerged as an excellent animal model of all types of human diseases (Bournele and Beis, 2016; Espín-Palazón et al., 2016; Hason and Bartůněk, 2019; Tyrkalska et al., 2016; Tyrkalska et al., 2019) due to its unique features, such as small size, large number of eggs generated at the same time, external fertilization, transparency, rapid development and easy and cheap maintenance (Diniz et al., 2015; Howe et al., 2013; Kimmel et al., 1995; Spence et al., 2008). In addition, zebrafish possesses high genetic and physiologic similarities with mammals, making it even more desirable (MacRae and Peterson, 2015). As an aquatic animal, zebrafish does not possess lungs, where silicosis develops, but does have a hindbrain ventricle, a cavity that is filled with cerebrospinal fluid into which immune cells can be recruited. This cavity can be used to mimic the alveolar environment in the human lungs and is a convenient injection site for silica crystals to study the molecular basis and pathogenesis of silicosis as well as for of drug screening (Cambier et al., 2014; Herbomel et al., 1999; Torraca et al., 2015).

Here, we establish for the first time a zebrafish model of silicosis that recapitulates the main characteristics of this disease, i.e. Nlrp3 inflammasome-driven inflammation and fibrosis. The combination therapy simultaneously targets the inflammasome and fibrogenesis, and shows enhanced efficacy compared to the monotherapy approach, opening new perspectives for targeted treatment of this devastating disease.

## RESULTS

### Injection of silica crystals into the hindbrain of zebrafish larvae promotes local and systemic inflammation

Here we present for the first time a zebrafish model of silicosis based on the injection of SiO_2_ crystals into the hindbrain ventricle of 48 hour-postfertilization (hpf) zebrafish larvae (Figure S1A). The endotoxin- and lipoprotein-free inorganic SiO_2_ crystals used in this study were very small in size, with a diameter of less than 100 nm, and with no characteristics that would make them useful for visualization *in vivo* in zebrafish. As most crystals, SiO_2_ crystals polarize light, but the *in vivo* site of the injection impeded the use of this feature, since bones and cartilages are also able to strongly polarize light, which made it impossible to distinguish between fish tissue and crystals *in vivo*. To ensure that SiO_2_ crystals left the needle during the microinjection procedures, we checked for the presence of crystals in the droplets from the needle on a slide by using their light polarization property (Figure S1B).

The presence of SiO_2_ crystals induced the recruitment to the injection site of fluorescent neutrophils and macrophages, which were labeled using the specific promoter myeloperoxidase (*mpx*) and microfibril-associated protein 4 (*mfap4*), respectively. As early as 1-hour post-injection (hpi) both neutrophils and macrophages were present in significant numbers in the hindbrain of the larvae injected with SiO_2_ crystals, while in the control larvae injected with PBS the number of both cell types was negligible (Figures 1A-1B and 2A-2B). These differences remained until 9 hpi in the case of neutrophils and 24 hpi in case of macrophages (Figures 1A-1B and 2A-2B), suggesting that SiO_2_ crystals can be recognized by the zebrafish immune system. Moreover, larvae injected with SiO_2_ crystals showed both neutrophilia and monocytosis at 24 hpi (Figures 1C-1D and 2C-2D), suggesting that SiO_2_ crystals activated emergency myelopoiesis. Interestingly, SiO_2_ crystals also increased neutrophil dispersion, assayed as the number of neutrophils outside the caudal hematopoietic tissues (CHT) at 24 hpi (Figure 1E-1F). Additionally, silica crystals were seen to have increased the number of macrophages in the head at 24 hpi (Figure 2E-2F). Interestingly, although SiO_2_ crystals injection did not affect larval survival (Figure S1C), they changed their behavior (Figure S1D). Thus, at 3 dpi zebrafish larvae injected with SiO_2_ crystals were less motile and remain in the middle of the Petri dish, in contrast to the control larvae that were very motile and always exploring the edges of the dish. Moreover, the escape response to a touch stimulus was impaired in silica-injected fish, and while control larvae instantly reacted by swimming rapidly away when touched on the end of the tail, SiO_2_-injected larvae showed either a very weak response or no response at all (Figure S1E).

**Figure 1:**
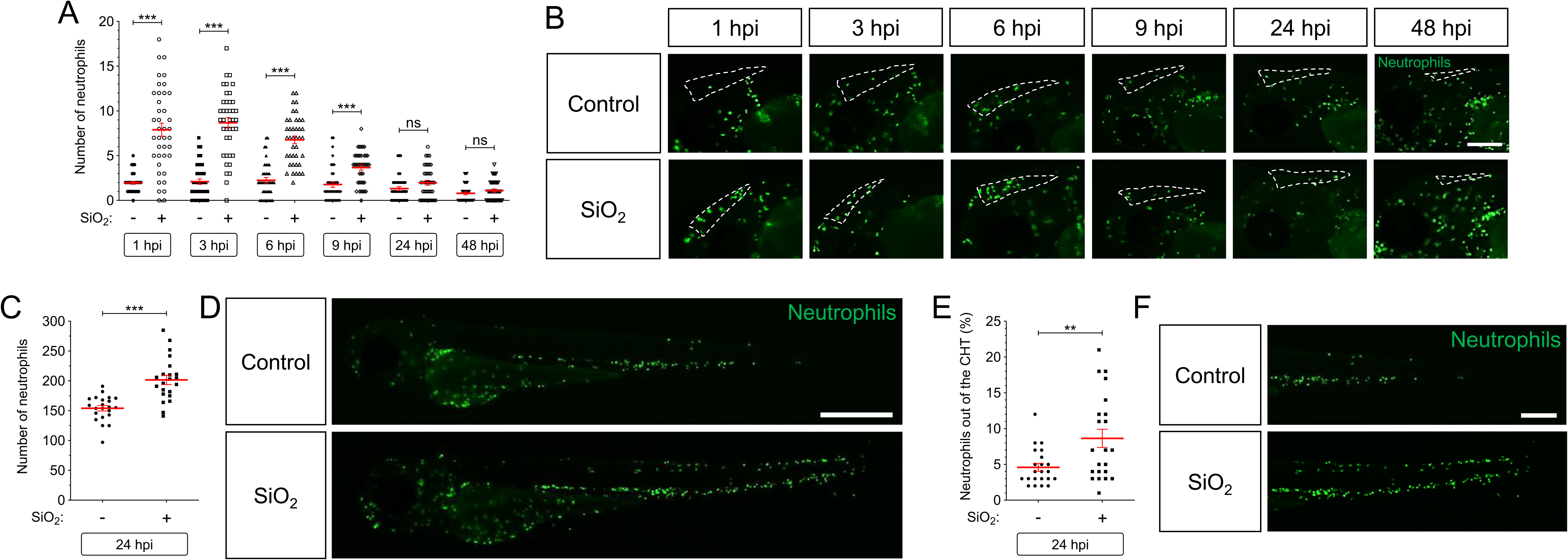
Silica crystals promotes neutrophil recruitment and neutrophilia in zebrafish larvae. SiO_2_ crystals or PBS were injected in the hindbrain ventricle (HBV, dash line) of 2 dpf *Tg(mpx:eGFP)*. Neutrophil recruitment (A, B), number in the whole body (C, D) and dispersion (E, F) were analyzed by fluorescence microscopy from 1 to 48 hpi (A, B) or 24 hpi (C-F). Each dot represents one individual and the mean ± S.E.M. for each group is also shown. P values were calculated using one-way ANOVA and Tukey multiple range test. ns, not significant, **p≤0.01, ***p≤0.001. Bars: 100 µm (B), 500 µm (D), 250 µm (F).

**Figure 2:**
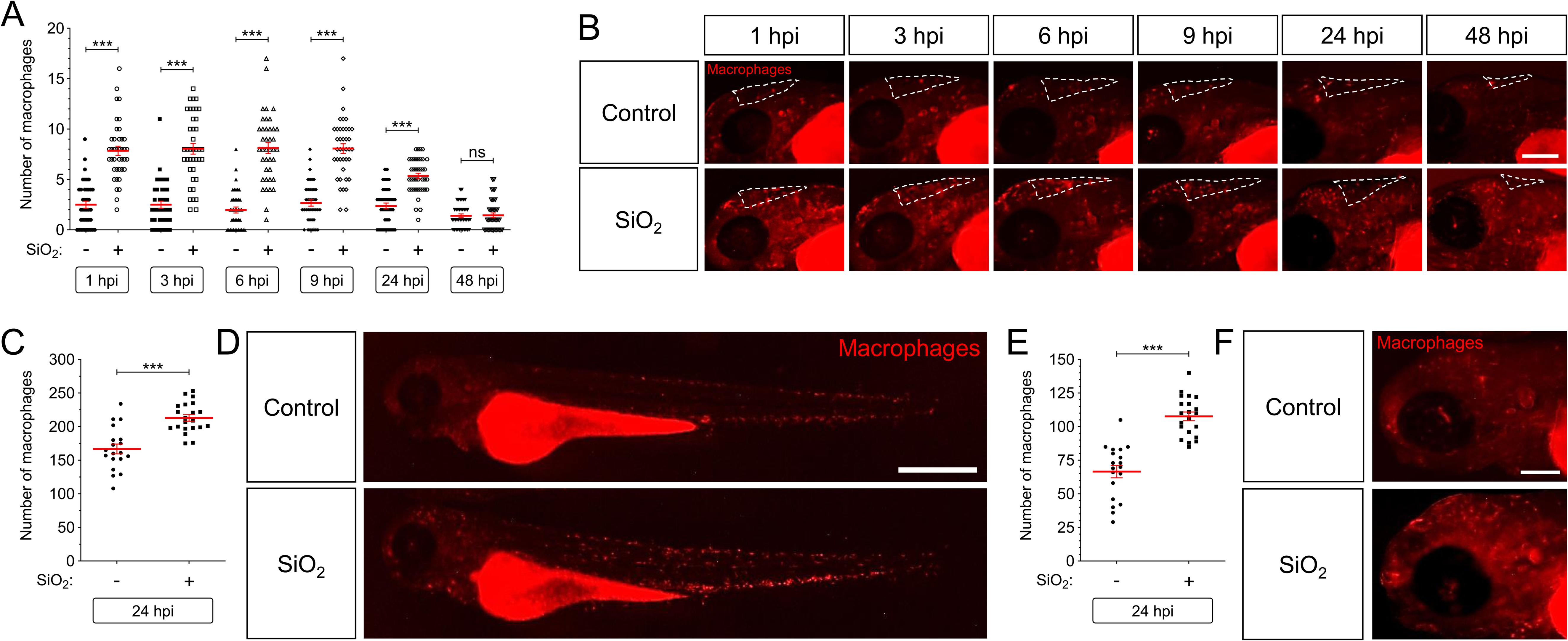
Silica crystals promotes macrophage recruitment and monocytosis in zebrafish larvae. SiO_2_ crystals or PBS were injected in the hindbrain ventricle (HBV, dash line) of 2 dpf *Tg(mfap4:mCherry)*. Macrophage recruitment (A, B) and number in the whole body (C, D) and the head (E, F) were analyzed by fluorescence microscopy from 1 to 48 hpi (A, B) or 24 hpi (C-F). Each dot represents one individual and the mean ± S.E.M. for each group is also shown. P values were calculated using one-way ANOVA and Tukey multiple range test. ns, not significant, ***p≤0.001. Bars: 100 µm (B), 500 µm (D), 100 µm (F).

To check the progress of the inflammatory process in the fish injected with SiO_2_ crystals, we analyzed the expression pattern of the master regulator of the inflammatory response nuclear factor κB using the reporter line *nfkb:eGFP*. At 6 and 9 hpi the larvae injected with SiO_2_ crystals showed a significantly increased fluorescent signal of NF-κB at the site of the injection (Figure 3A-3B). The NF-κB not only increased locally, but also robustly increased systemically at 24 hpi (Figure 3C-3D). The expression pattern of the gene encoding the main proinflammatory cytokine interleukin-1β (Il1b) was then analyzed using the reporter line *il1b:eGFP*. At 6, 9 and 24 hpi, the fluorescent signal levels at the site of the injection had significantly increased in the larvae injected with SiO_2_ crystals (Figure 3E-3F). In contrast to *nfkb*, *il1b* levels only increased locally at the site of the injection (Figure 3G-3H). Finally, the expression pattern of the gene encoding tumor necrosis factor α (Tnfa), a proinflammatory cytokine mostly produced by macrophages, was analyzed using the reporter line *tnfa:eGFP*. By 3 hpi the fluorescent signal at the site of the injection had significantly increased and remained high until 9 hpi, the cells producing Tnfa showing a clearly marked outline (Figure S2A-S2B). As in the case of *il1b*, *tnfa* levels only increased locally at the site of the injection (Figure S2C-S2D). To confirm that the cells responsible for Tnfa production were macrophages, we used a zebrafish line in which macrophages were labeled in red using the macrophage-expressed gene 1 (*mpeg1*) promoter (*mpeg1:cherry*) and cells producing Tnfa in green (*tnfa:eGFP*). At 6 hpi more than 40% of macrophages of larvae injected with SiO_2_ crystals were seen to be producing Tnfa, in contrast to the negligible amount of Tnfa^+^ macrophages observed in PBS-injected larvae (Figure S2E and S2G). Most of the Tnfa^+^ macrophages were in the head of the SiO_2_ crystal-injected larvae, where they represented more than 50% of macrophages (Figure S2F-S2G).

**Figure 3:**
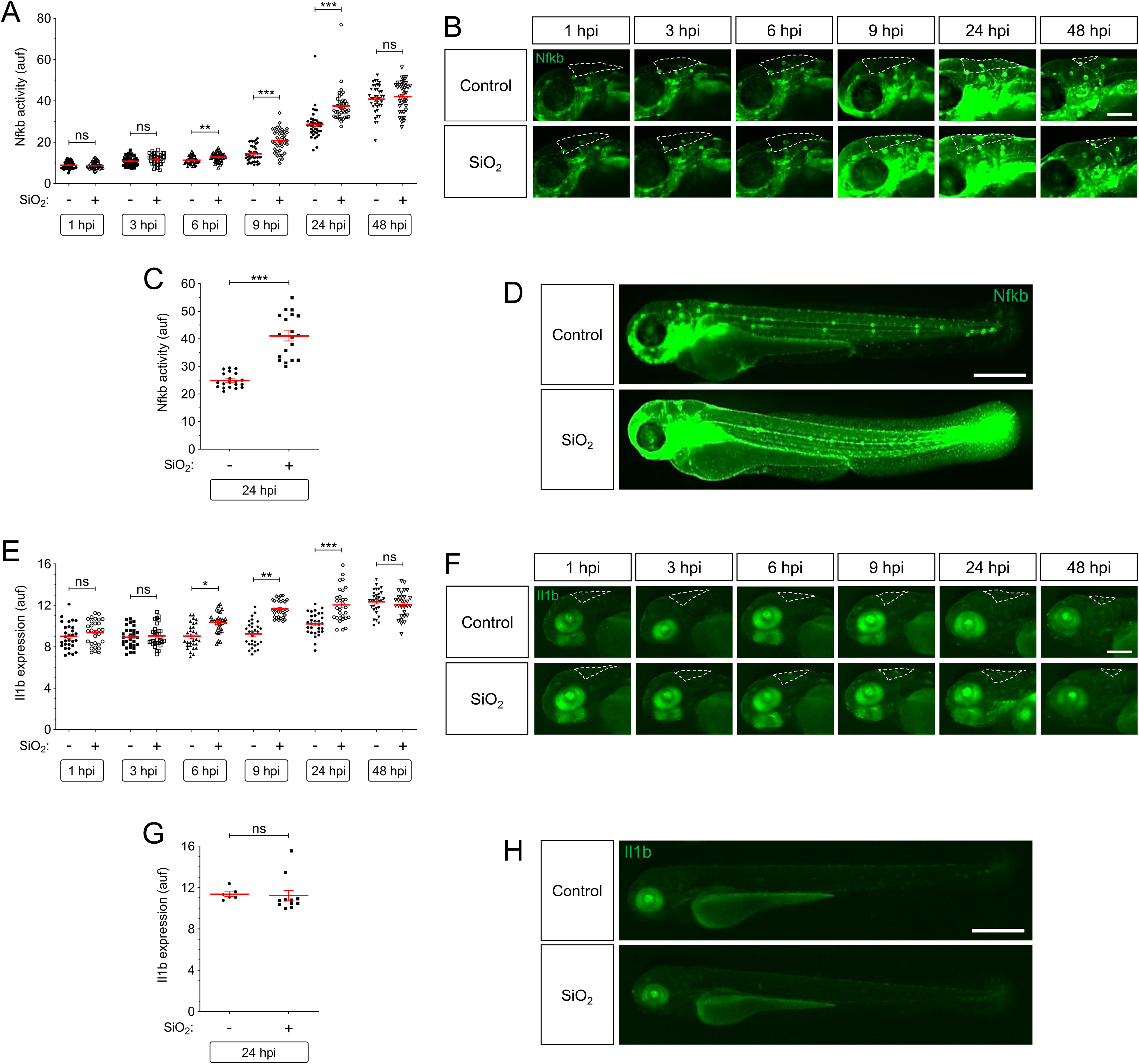
Silica crystals promotes local and systemic inflammation in zebrafish larvae. SiO_2_ crystals or PBS were injected in the hindbrain ventricle (HBV, dash line) of 2 dpf *Tg(NFkB-RE:eGFP)* (A-D) and *Tg(il1b:eGFP)* (E-H). Nfkb activation (A-D) and il1b production (E-H) were analyzed by fluorescence microscopy from 1 to 48 hpi (A, B, E, F) or 24 hpi (C, D, G, H). Each dot represents one individual and the mean ± S.E.M. for each group is also shown. P values were calculated using one-way ANOVA and Tukey multiple range test. ns, not significant, *p≤0.05, **p≤0.01, ***p≤0.001. Bars: 100 µm (B, F), 500 µm (D, H).

The induction of genes encoding inflammatory mediators at the injection site at 6 and 12 hpi was further confirmed using RT-qPCR. The expression levels of *il1b*, *nfkb1*, *cxcl8a* (chemokine CXC motif ligand 8a), *tnfa*, *ptgs2b* (prostaglandin-endoperoxide synthase 2b) and *il10* had significantly increased in larvae injected with SiO_2_ crystals at both time points, whereas *ptgs2a* was only seen to have increased significantly at 12 hpi (Figures S3A-S3G). A similar trend was observed in the transcript levels of genes encoding the main inflammasome components, Caspa (caspase a), Asc (*pycard* gene) and Nlrp3 (NLR family pyrin domain containing 3) (Figures S3H-S3J), whereas those of the gene encoding its paralog Nlrp3l (Nlrp3-like) did not increase following SiO_2_ crystal injection (Figure S3K). Interestingly, caspase-1 activity only increased systemically in SiO_2_ crystal-injected larvae (Figure S3L). PBS injection did not result in an inflammatory response or significant immune cell recruitment to the site of the injection in any of the experiments (Figures 1-3 and S2-S3). Collectively, these results suggest that SiO_2_ crystals were able to induce both local and systemic inflammation in zebrafish.

### The Cxcl8/Cxcr2 axis is required for the spreading of silica crystal-induced inflammation

CXCL8 (C-X-C Motif Chemokine Ligand 8, also known as IL-8), a cytokine secreted by endothelial, epithelial and immune cells (Russo et al., 2014), plays an important role as a major mediator of neutrophil recruitment via engagement of the two G protein-coupled receptors CXCR1 and CXCR2 in mammals (Ha et al., 2017) and zebrafish (de Oliveira et al., 2015; Gabellini et al., 2018). To check whether myeloid cells need to be recruited for the inflammatory response by silica crystals to be induced, zebrafish larvae were treated with the specific CXCR2 inhibitor SB225002, which is able to inhibit neutrophil recruitment in zebrafish (de Oliveira et al., 2015). The results showed that the number of neutrophils and macrophages recruited to the hindbrain decreased in SB225002-treated larvae after SiO_2_ injection at all analyzed timepoints (Figure 4A and 4B). Strikingly, the pharmacological inhibition of Cxcr2 blocked neutrophilia and monocytosis in larvae injected with SiO_2_ crystals, without affecting the basal numbers of neutrophils and macrophages (Figure 4A and 4B). Interestingly, while the inhibition of Cxcr2 also impaired the systemic induction of *il1b*, *cxcl8a*, *nfkb1*, *tnfa* and *il10* transcript levels by SiO_2_ crystals, they were largely unaffected at the injection site (Figure 4C-4G). In contrast, *ptgs2a* and *ptgs2b* mRNA levels increased locally and systemically upon SB225002 treatment in SiO_2_ crystal-injected larvae (Figure 4G-4H). In addition, the genes encoding inflammasome components were largely unaffected by inhibition of Cxcr2 in SiO_2_ crystal-injected larvae (Figures 4J-4M). These results suggest that the Cxcl8/Cxcr2 axis is instrumental for the spreading of the inflammatory response induced by silica crystals.

**Figure 4:**
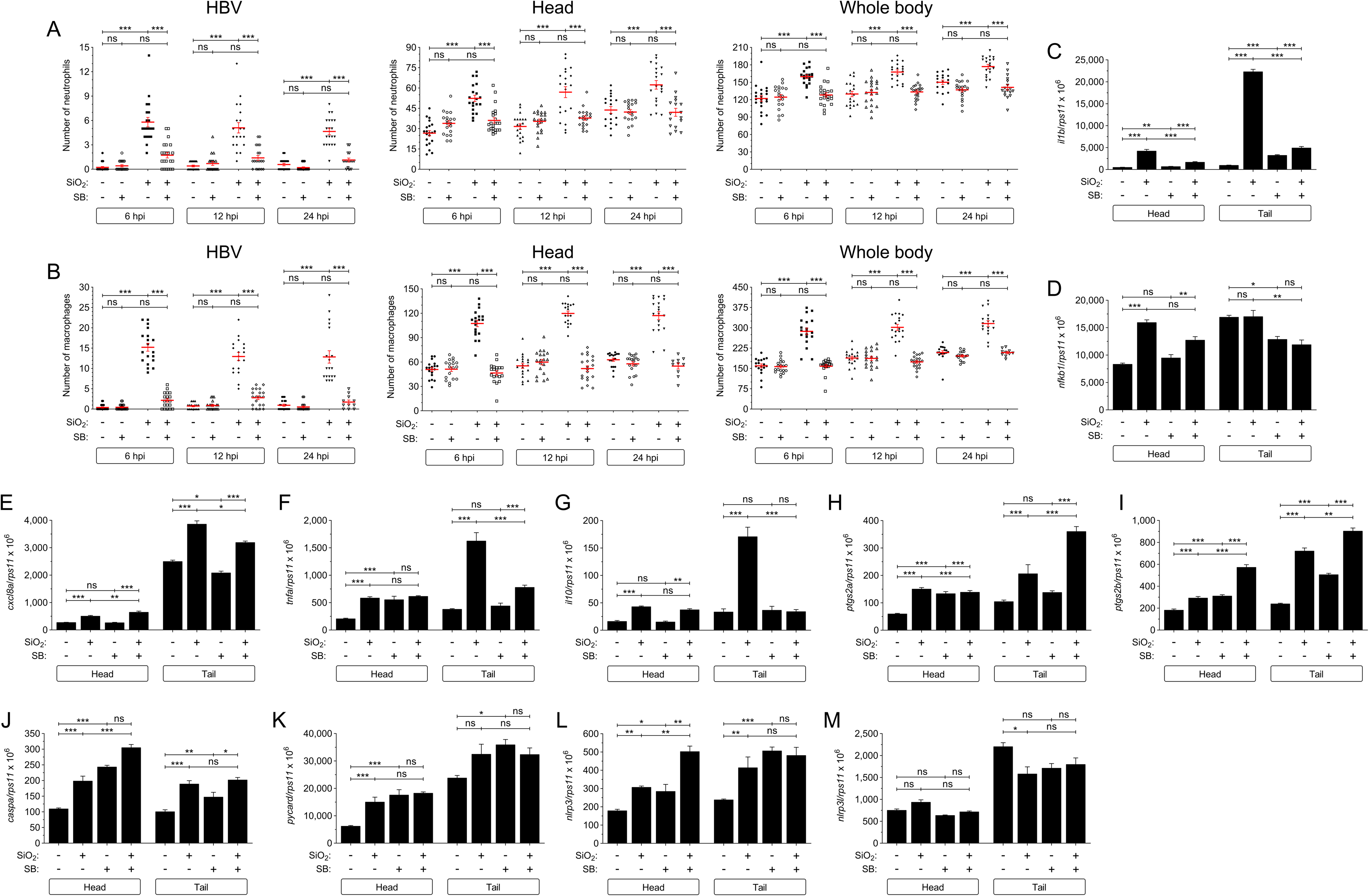
The Cxcl8/Cxcr2 axis is required for the spreading of silica crystal-induced inflammation. SiO_2_ crystals or PBS were injected in the hindbrain ventricle (HBV) of 2 dpf *Tg(lyz:DsRED2)* (A), *Tg(mfap4:mCherry)* (B) and wild type (C-M) larvae in the presence of either DMSO or the Cxcr2 inhibitor SB225002 (SB). Neutrophil (A) and macrophage (B) recruitment and number were analyzed at 6, 12 and 24 hpi by fluorescence microscopy, while the transcript levels of the indicated genes (C-M) were analyzed at 12 hpi by RT-qPCR in larval head and tail. Each dot represents one individual and the mean ± S.E.M. for each group is also shown. P values were calculated using one-way ANOVA and Tukey multiple range test. ns, not significant, *≤p0.05, **p≤0.01, ***p≤0.001.

### Silica crystals signal through the Nlrp3/Caspa/Gsdme inflammasome and Myd88 in zebrafish

It has been shown that the NLRP3 inflammasome plays an important role in the development and progression of silicosis (Cassel et al., 2008; Dostert et al., 2008; Peeters et al., 2014). To ascertain whether the inflammasome mediates the proinflammatory activity of SiO_2_ crystals in zebrafish, larvae were treated with the specific caspase-1 inhibitor VX-765 by bath immersion. Interestingly, Although the pharmacological inhibition of caspase-1 failed to impair the recruitment of neutrophils to the hindbrain, it was able to fully rescue the neutrophilia induced by SiO_2_ crystals (Figure 5A). In addition, pharmacological inhibition of the canonical inflammasome also robustly decreased the induction of the inflammatory genes caused by the injection of SiO_2_ crystals into the hindbrain of the zebrafish larvae (Figures 5B-5I). Moreover, *caspa* and *nlrp3* transcript levels were strongly induced when SiO_2_ crystal-injected larvae were treated with VX-765 (Figure 5I and 5L), while *pycard* and *nlrp3l* levels were unchanged (Figure 5J and 5K). As expected, VX-765 decreased caspase-1 activity in both PBS- and SiO_2_ crystal-injected larvae (Figure 5M). All these results indicate that silica crystals activate the canonical inflammasome in zebrafish, leading to inflammation and emergency myelopoiesis.

**Figure 5:**
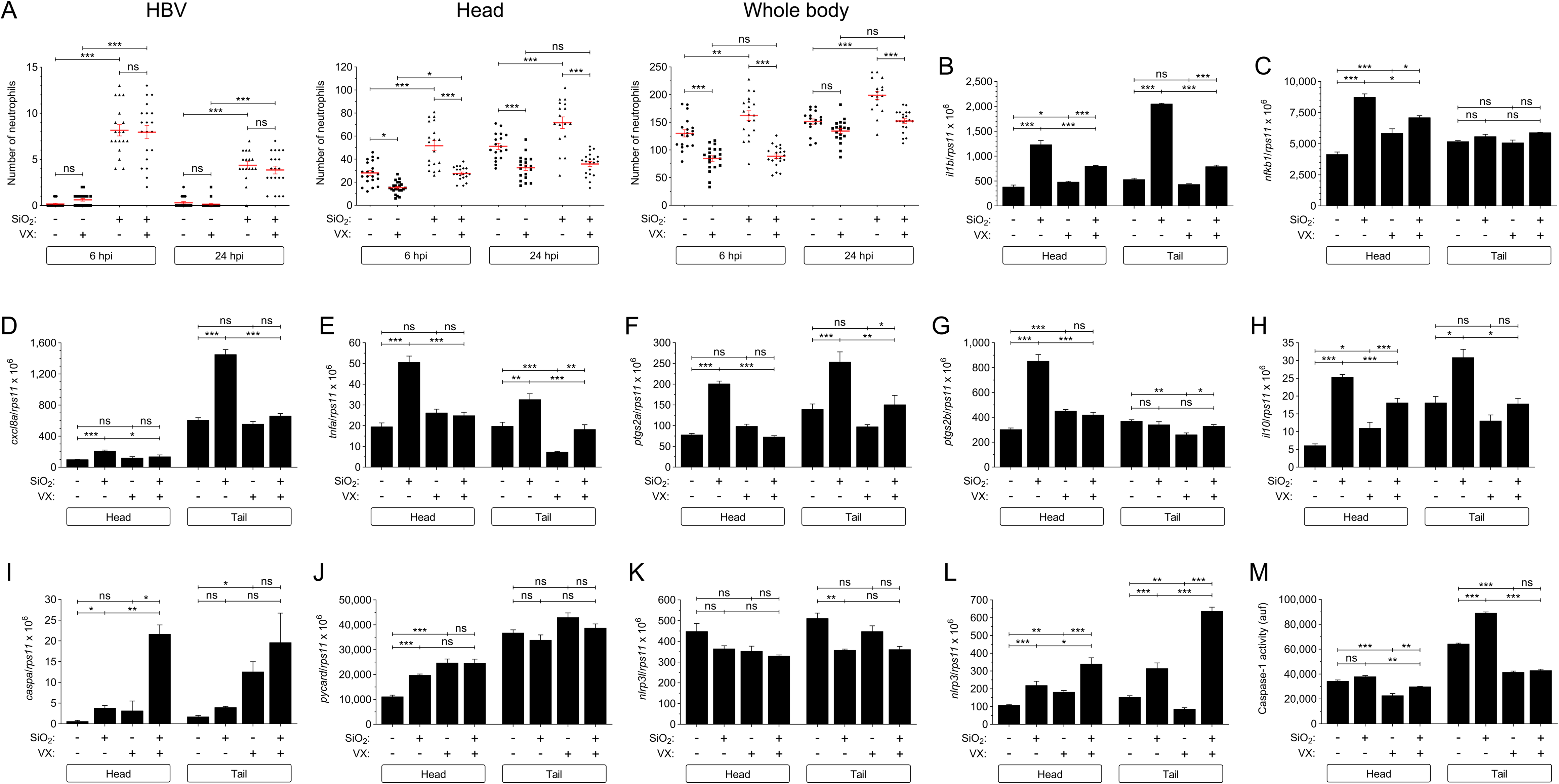
Silica crystals signal through the canonical inflammasome in zebrafish. SiO_2_ crystals or PBS were injected in the hindbrain ventricle (HBV) of 2 dpf *Tg(lyz:DsRED2)* (A) and wild type (B-M) larvae in the presence of either DMSO or the caspase-1 inhibitor VX-765 (VX). Neutrophil (A) recruitment and number were analyzed at 6, 12 and 24 hpi by fluorescence microscopy, while the transcript levels of the indicated genes (B-L) were analyzed at 12 hpi by RT-qPCR in larval head and tail. Caspase-1 activity levels were analyzed at 24 hpi in larval head and tail (M). Each dot represents one individual and the mean ± S.E.M. for each group is also shown. P values were calculated using one-way ANOVA and Tukey multiple range test. ns, not significant, *≤p0.05, **p≤0.01, ***p≤0.001.

Gasdermins are the executors of programmed cell death known as pyroptosis and they play a crucial role in infection and inflammatory diseases (Broz et al., 2020). It has been reported that zebrafish possess 2 paralogs of Gsdme, called Gsdmea and Gsdmeb, that can be cleaved by zebrafish caspase-1 (Chen et al., 2021) and regulate neutrophil and macrophage pyroptosis (Lozano-Gil et al., 2022). We found that genetic inhibition of Gsdmea/b with CRISPR/Cas9 (about 60% knockout efficiency for both genes) resulted in increased neutrophil and macrophage recruitment by SiO_2_ crystals, and enhanced neutrophilia and monocytosis at all time points analyzed (Figure 6A and 6B), suggesting the inhibition of neutrophil and macrophage pyroptotic cell death (Lozano-Gil et al., 2022). Moreover, Gsdmea/b deficiency led to increased Nfkb levels at the site of the injection, and in the head and the whole body in PBS- and SiO_2_ crystal-injected larvae at all analyzed timepoints, although the induction was significantly higher in the larvae injected with SiO_2_ crystals (Figure 6C). These results were further confirmed by the higher transcript levels of genes encoding inflammatory mediators and inflammasome components found in SiO_2_ crystal-injected larvae than in the corresponding controls (Figure S4A-S4J), apart from *nlrp3l* gene (Figure S4K). As expected, no difference in caspase-1 activity was observed between control and Gsdmea/b-deficient larvae (Figure S4L). However, the inhibition of caspase-1 activity by VX-765 rescued SiO_2_ crystal-induced neutrophilia in Gsdmea/b-deficient larvae at 6, 12 and 24 hpi (Figure 6D). Taken together, these results suggest that silica crystals activate two signaling pathways: a Caspa/Gsdmea/b axis that regulates myeloid cell pyroptosis and inflammation (Lozano-Gil et al., 2022), and a Caspa/Gata1 axis that promotes emergency myelopoesis (Tyrkalska et al., 2019).

**Figure 6:**
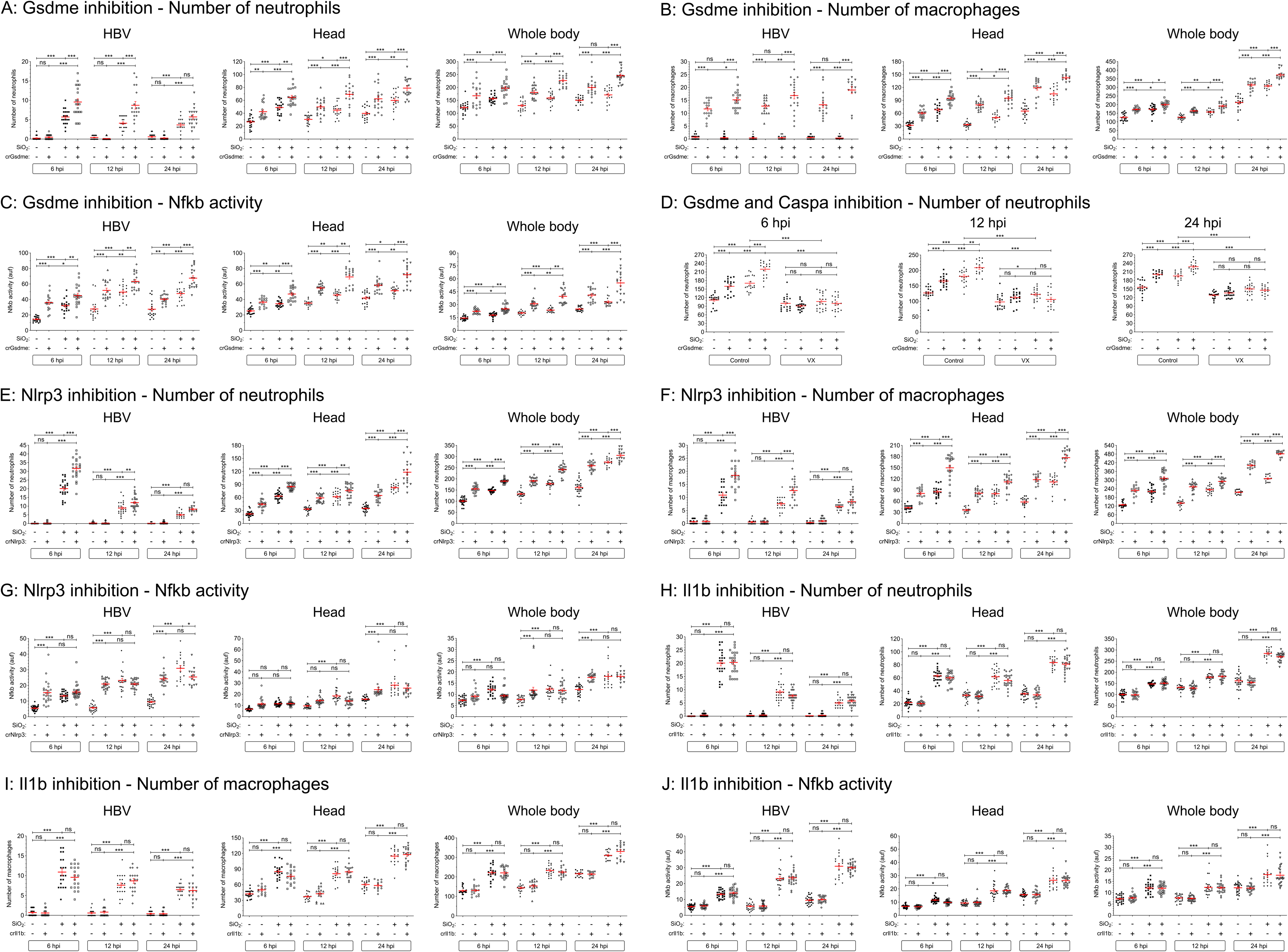
Silica crystals signal through the Nlrp3/Caspa/Gsdme inflammasome in zebrafish. *Tg(lyz:DsRED2)* (A, D, E, H), *Tg(mfap4:mCherry)* (B, F, I) and *Tg(NFkB-RE:eGFP)* (C, G, J) 1-8 cell stage embryos were microinjected with control, *gsdmea/b* (A-D), *nlrp3* (E-G) or *il1b* (H-J) crRNA/Cas9 complexes. At 2 dpf, SiO_2_ crystals or PBS were injected in the larval hindbrain ventricle (HBV) in the presence of either DMSO or the caspase-1 inhibitor VX-765 (VX) (D). Neutrophil (A, D, E, H) and macrophage (B, F, I) recruitment and number, and Nfkb activity (C, G, J) were analyzed at 6, 12 and 24 hpi by fluorescence microscopy. Each dot represents one individual and the mean ± S.E.M. for each group is also shown. P values were calculated using one-way ANOVA and Tukey multiple range test. ns, not significant, *≤p0.05, **p≤0.01, ***p≤0.001.

NLRP3 inflammasome is already known to take part in the silica crystal response in patients (Peeters et al., 2014), where the stimulation of macrophages with silica crystals results in the activation of caspase-1 in a NLRP3-dependent manner (Cassel et al., 2008). Nlrp3 deficiency (knockout efficiency of 75%) resulted in increased SiO_2_ crystal-induced neutrophil and macrophage recruitment, and neutrophilia and monocytosis at 6, 12 and 24 hpi (Figure 6E and 6F), phenocopying Gsdmea/b deficiency. Moreover, although SiO_2_ crystals were able to increase Nfkb to similar levels in both PBS- and SiO_2_ crystal-injected larvae (Figure 6G), the transcript levels of genes encoding inflammatory mediators and inflammasome components (Figure S5A-S5) were weakly attenuated in Nlrp3-deficient larvae. However, the genes encoding Il10 (Figure S5G) and inflammasome components were hardly affected, apart from the gene encoding Nlrp3 itself (Figure S5H-S5K). As expected, caspase-1 activity was significantly lower in Nlrp3-deficient larvae injected with either PBS or SiO_2_ (Figure S5I).

To check whether the activation of the inflammasome by silica crystals promoted inflammation through the cleavage of Il1b, this gene was inactivated using CRISPR/Cas9 technology (knockout efficiency of 75%) before checking neutrophil and macrophage recruitment to SiO_2_ crystals, Nfkb activation and gene expression. Il1b deficiency had no effect on neutrophil and macrophage recruitment to SiO_2_ crystals, and neutrophilia and monocytosis at 6, 12 and 24 hpi (Figures 6H and 6I). Similarly, Nfkb levels were similar at the site of the injection, head, and the whole body in PBS- and SiO_2_ crystal-injected larvae at all the analyzed timepoints (Figure 6J). These results were further confirmed by analyzing the transcript levels of genes encoding inflammatory mediators and inflammasome components that were similarly induced in wild type and Il1b-deficient larvae by SiO_2_ crystals, except those of *il1b* that were higher in the mutant than in wild type larvae (Figure S6A-S6K), probably reflecting a compensatory mechanism.

It was then checked whether silica crystals also signal via Myd88, the main adaptor protein required for TLR signaling, using a Myd88-deficient zebrafish line (van der Vaart et al., 2013). Although the injection of SiO_2_ crystals in the hindbrain of Myd88-deficient larvae failed to increase the transcript levels of genes encoding inflammatory mediators (Figures 7A-7F), the levels of genes encoding inflammasome components were induced at similar levels in wild type and Myd88-deficient larvae (Figures 7H-7K), suggesting that silica crystals activate both TLR and inflammasome signaling pathways in zebrafish.

**Figure 7:**
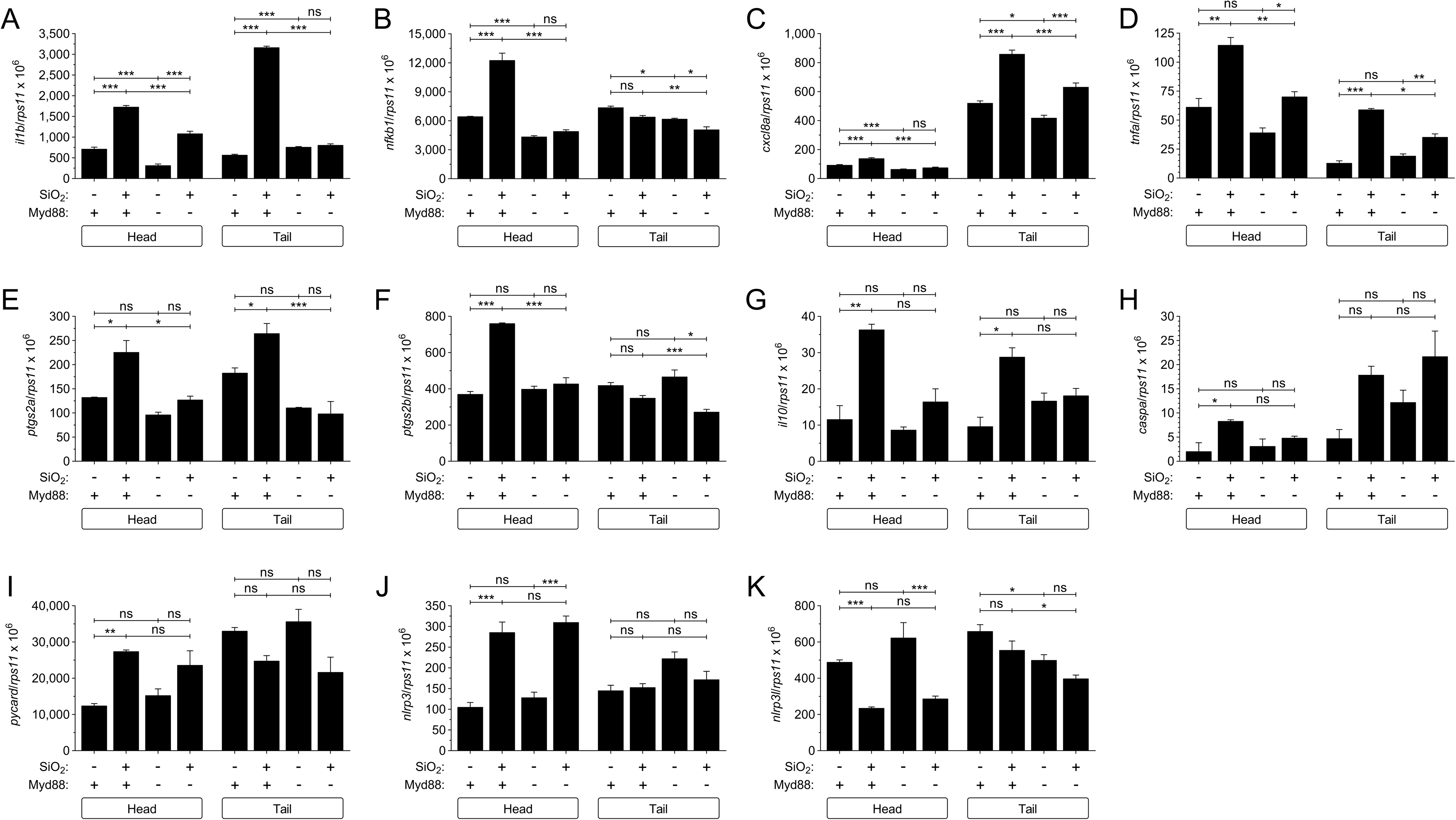
Silica crystals signal through Myd88 in zebrafish. SiO_2_ crystals or PBS were injected in the hindbrain ventricle (HBV) of 2 dpf wild type and Myd88-deficient larvae. The transcript levels of the indicated genes were analyzed at 12 hpi by RT-qPCR in larval head and tail. The data are shown as the mean ± S.E.M. P values were calculated using one-way ANOVA and Tukey multiple range test. ns, not significant, *≤p0.05, **p≤0.01, ***p≤0.001.

### Silica crystals promote fibrosis in zebrafish

The endocytosis of silica crystals by macrophages initiates the expression of transforming growth factor-β1 (TGF-β1), which promotes the transcription of extracellular matrix gene expression in lung fibroblasts (Allison, 2014; Davis et al., 1996). Therefore, we checked the transcript levels of *tgfb1* at 6, 12, 24, 72 and 120 hpi, finding that SiO_2_ crystal injection resulted in increased transcript levels of *tgfb1* at 12, 24, 48 and 120 hpi (Figure 8A). A similar pattern was seen for *acta2*, which encodes the pro-fibrotic zebrafish biomarker smooth muscle actin alpha 2 (van der Helm et al., 2018), and for *col1a1a*, *col1a2* and *fn1a*, which encodes type I collagens alpha 1a and alpha 2, and fibronectin 1a, respectively (Figures 8B-E). However, *acta2*, *col1a2* and *fn1a* transcript levels were induced by SiO_2_ crystals not only at the injection site, but also systemically (Figures 8B, 8D and 8E). In contrast, *il1b* mRNA levels returned to basal levels by 24 hpi (Figure 8F), suggesting that fibrosis started after the resolution of inflammation.

**Figure 8:**
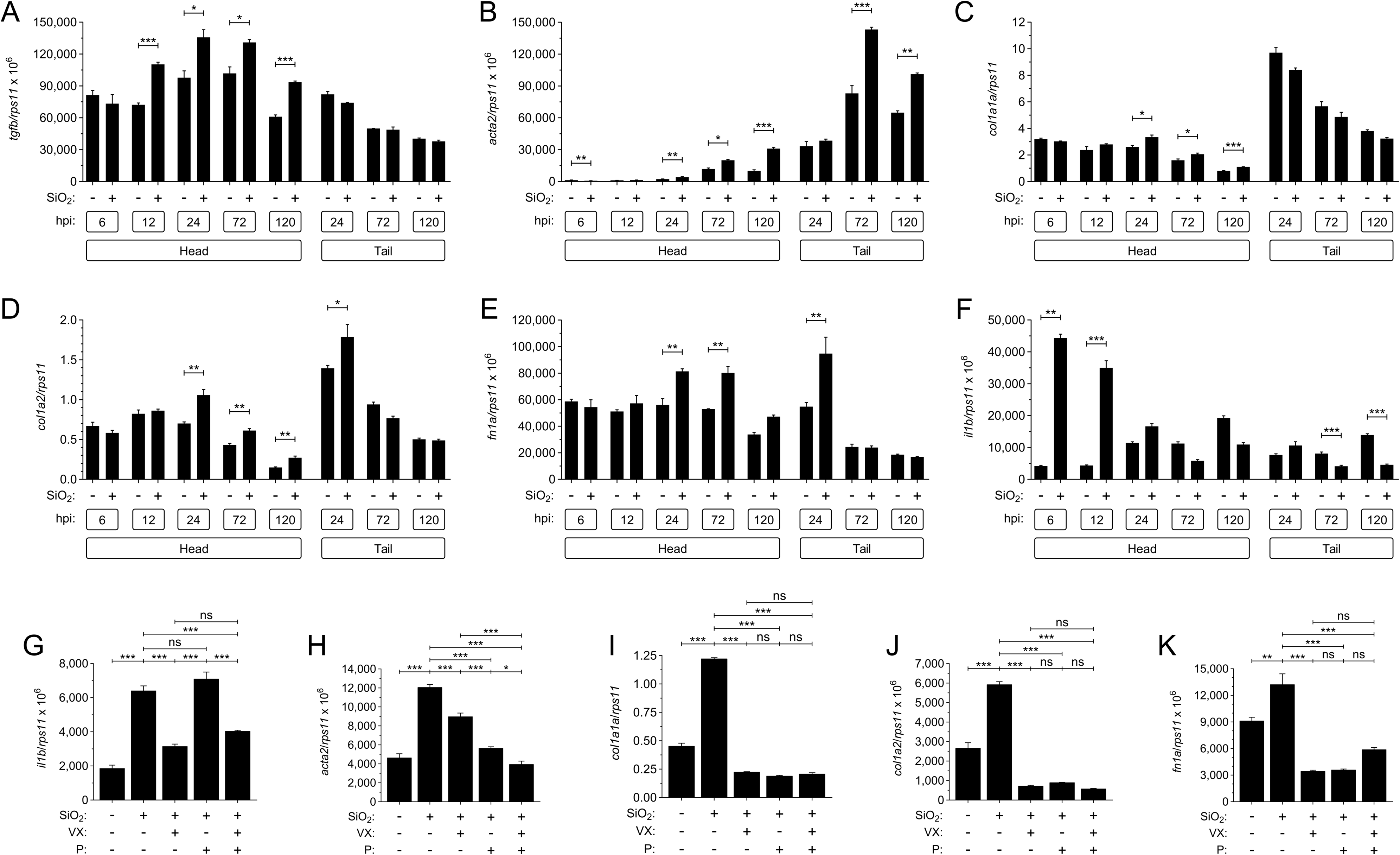
Cotreatment with VX-765 and pirfenidone alleviates inflammation and fibrosis induced by silica crystals in zebrafish larvae. SiO_2_ crystals or PBS were injected in the hindbrain ventricle (HBV) of 2 dpf wild type larvae alone (A-F), or in the presence of DMSO or the combination of the caspase-1 inhibitor VX-765 (VX) and/or the antifibrotic drug pirfenidone (P) (G-K). The transcript levels of the indicated genes were analyzed from 6 to 120 hpi in larval head and tail (A-F) or 120 hpi in head (G-K) by RT-qPCR. The data are shown as the mean ± S.E.M. P values were calculated using one-way ANOVA and Tukey multiple range test. ns, not significant, *≤p0.05, **p≤0.01, ***p≤0.001.

### Cotreatment with VX-765 and pirfenidone alleviates inflammation and fibrosis induced by silica crystals in zebrafish larvae

As the therapeutic options for silicosis patients are limited, we decided to check the efficiency of the caspase-1 inhibitor VX-765 and the antifibrotic drugs nintedanib, pirfenidone and halofuginone hydrobromide alone or in combination. Firstly, it was checked that the drugs were not toxic at the working concentrations during chronic exposure during larval development (5 days of incubation) (Figure S7A-S7C). It was observed that although VX-765 was unable to decrease *acta2* and *fn1a* transcript levels, it decreased those of *col1a1a* and *col1a2* (Figure S8A-S8D). In contrast, pirfenidone and halofuginone hydrobromide decreased the transcript levels of *acta2*, *col1a1a*, *col1a2* and *fn1a* in the heads of the larvae injected with SiO_2_ crystals (Figure S8E-L). However, although nintedanib decreased the mRNA levels of *acta2*, *col1a1a*, *col1a2*, it paradoxically increased those of *fn1a* (Figure S8M-S8P). Finally, as none of the treatments was able to restore inflammation and fibrotic markers to basal levels in SiO_2_ crystal-injected larvae, we tried the combination of VX-765 and pirfenidone to target both inflammation and fibrinogenesis. Strikingly, this cotreatment robustly decreased inflammation and fibrosis in SiO_2_ crystal-injected larvae, since the transcript levels of all inflammatory and fibrotic markers analyzed, i.e. *acta2*, *fn1*, *col1a1a*, *col1a2* and *il1b*, fell to basal levels and were lower than in larvae treated with either drug alone (Figures S8F-8K).

## DISCUSSION

Although mouse and rat models of silicosis have provided some mechanistic insight this disease, there is controversy about the signaling pathways involved and, more importantly, the therapeutic options for patients are rather limited. We describe a zebrafish model of silicosis that provides novel data about the mechanisms orchestrating this pathology and constitute a unique platform for chemical screening. Our data revealed that both TLR and inflammasome signaling pathways cooperate to drive silica crystals-induced inflammation and fibrosis. Although the relevance of TLRs in lung fibrosis has been established in mouse models (Li et al., 2015) and confirmed in epidemiological studies in human patients (McElroy et al., 2022), its involvement in silicosis is controversial. While a TLR4/MyD88 pathway has been shown to be involved in silica-induced inflammation in U937-differentiated macrophages (Chan et al., 2018), silica crystals attenuate TLR ligand-dependent dendritic cell activation (Beamer and Holian, 2008). In addition, intranasal instillation of TMX-306, a weak agonist of TLR7, reduced silica crystal-induced pulmonary granuloma, fibrotic lesions, TGF-β production and, surprisingly, silica particles present in the interstitial space (Ferreira et al., 2016). Our data support a key role for TLR signaling in the induction of inflammation by silica crystals, since Myd88-deficient fish were highly resistant to silica crystal-induced inflammation, whereas their Il1b-deficient counterparts responded in the same way as wild type animals, ruling out the impact of Myd88 deficiency in the zebrafish model of silicosis was via Il1b signaling. This result was unexpected, since neutralization of IL-1β with the IL-1 type I receptor (IL-1R) antagonist (Song et al., 2014) and IL-1R deficiency (Hornung et al., 2008) reduce silica inflammation in mouse models of silicosis. Furthermore, the compassionate use of anakinra in a miner diagnosed with silicosis led to a gradual improvement in respiratory symptoms and normalization of the inflammatory markers, such as C reactive protein and erythrocyte sedimentation (Cavalli et al., 2015). However, randomized clinical trials with appropriate placebo group are required to definitively confirm the efficacy of anakinra and the involvement of IL-1 β in this much overlooked disease.

Two seminal studies demonstrate that silica crystals activate the NLRP3 inflammasome in mouse macrophages and human peripheral blood mononuclear cells, leading to the release of IL-1β and IL-18 (Cassel et al., 2008; Hornung et al., 2008). Similarly, an Nlrp3/Caspa/Gsdme inflammasome is activated and contributes to the inflammatory and fibrotic processes induced by silica crystals. Thus, pharmacological inhibition of caspase-1 activity and the genetic inhibition of Nlrp3 alleviated inflammation and fibrosis in this model, although they had an opposite effect on neutrophilia and monocytosis. In sharp contrast, genetic inhibition of Gsdmea/b promoted hyperinflammation and increased neutrophilia and monocytosis in response to silica crystals, all these effects being dependent on caspase-1 activity. These unexpected results could be reconciled if we consider the involvement of two inflammasomes (Figure 9A): (1) inhibition of caspase-1 activity, i.e. Caspa, blocks not only the Nlrp3 inflammasome, but also, the inflammasome responsible for the cleavage of Gata1 (Tyrkalska et al., 2019), resulting in impaired basal and emergency myelopoiesis, and reduced inflammation; (ii) inhibition of Nlrp3 blocks pyroptotic cell death of neutrophils and macrophages (Lozano-Gil et al., 2022), which reduces inflammation despite the increased number of neutrophils and macrophages; and (iii) inhibition of Gsdme only impairs pyroptosis of myeloid cells leaving a Gsdme-independent inflammatory pathway unaffected. Whatever the outcome, our results point to the key role of emergency myelopoiesis and pyroptosis in silicosis, as occurs in other chronic inflammatory diseases (Newton et al., 2021). Although the relevance of pyroptosis has not been formally examined in mouse models of silicosis or patients, the exposure of primary human respiratory epithelial cells to silica particles induces pyroptosis (Chen et al., 2022). Nevertheless, cell death, assayed as lactate dehydrogenase release, induced by silica crystals is equivalent in wild type and NLRP3-deficient macrophages (Cassel et al., 2008). Therefore, further studies in animal models aimed at revealing the role played by pyroptotic cell death in silicosis are required.

**Figure 9:**
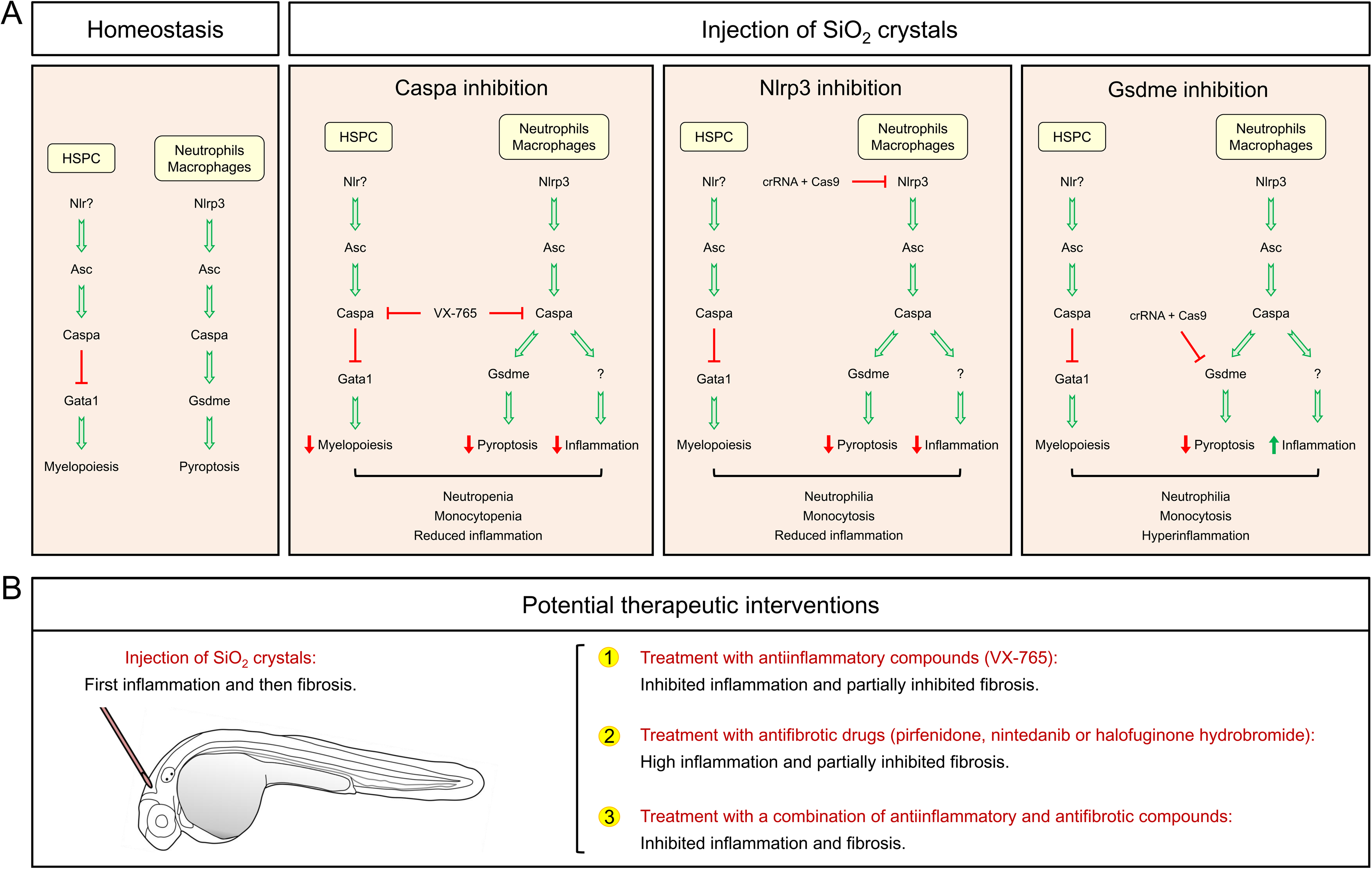
Working models. (A) Proposed model illustrating the impact of chemical and genetic inhibition of different inflammasome components in silica crystal-induced inflammation and emergency myelopoiesis. (B) Proposed therapeutic interventions for silicosis based in individual treatments targeting either inflammasome or fibrosis and their combination according to the results obtained in the zebrafish silicosis model developed in this study.

While it has been reported that neutrophils play a critical role in silicosis, as occurs in other inflammatory diseases, the relevance of monocyte infiltration is largely unknown. Inhibition of LTB4-dependent neutrophil infiltration reduces neutrophilic inflammation, and the depletion of circulating monocytes and neutrophils by intraperitoneal injection of cyclophosphamide reduces the influx of macrophages and neutrophils in the bronchoalveolar lavage in rodent models of silicosis (Hegde et al., 2018; Nemmar et al., 2005). The zebrafish silicosis model established here revealed the Cxcl8a/Cxcr2-dependent recruitment of both neutrophils and macrophages to silica crystals and, more importantly, their requirement to promote emergency myelopoiesis and spread inflammation. Although further investigation is required to ascertain whether myeloid cell recruitment and/or the Cxcl8a/Cxcr2 axis itself are mediating these effects, the almost unaltered local inflammation upon inhibition of Cxcr2 signaling strongly suggests that recruited myeloid cells are indispensable to spread inflammation and promote emergency myelopoiesis. These results are consistent with the reduced peripheral thrombogenicity observed in hamster model of silicosis upon selective ablation of circulating neutrophils and monocytes (Nemmar et al., 2005) and, importantly, with the obvious link between exposure to silica crystals and systemic autoimmune diseases revealed by epidemiological studies (Pollard, 2016).

One of the most interesting observations of our study is that, while single pharmacological inhibition of either the inflammasome with the caspase-1 inhibitor VX-765 or fibrosis with drugs targeting different signaling pathways, including VEGFR1/2/3, FGFR1/2/3 and PDGFRα/β with nintedanib, chemokine and collagen production in fibroblasts with pirfenidone, and type-I collagen synthesis with halofuginone hydrobromide, partially alleviated inflammation and fibrosis induced by silica crystals in zebrafish. However, the combination of VX-765 and pirfenidone robustly decreased silica crystal-induced inflammation and fibrosis (Figure 9B). Pirfenidone has also been shown to ameliorate silica inflammation and fibrosis in rodent models of silicosis (Cao et al., 2022; Guo et al., 2019). As pirferidone is approved for the treatment of idiopathic pulmonary fibrosis and a clinical trial with VX-765 has shown good tolerance and pharmacokinetic properties in patients (EudraCT Numbers 2004-000283-27 and 2011-004156-19), the combination therapy used here is seems to show promise for treating silicosis and other devastating pneumoconiosis.

In summary, we have developed a zebrafish model of silicosis that confirms the critical role played by the Nlpr3 inflammasome and TLR signaling in the inflammation and fibrosis induced by silica crystals *in vivo.* The model also shows for the first time the contribution of emergency myelopoiesis and myeloid cell-mediated spreading of inflammation in this disease. This model identifies an effective combination therapy that simultaneously targets inflammation and fibrosis, pointing to the potential for drug repurposing via highthroughput chemical screening.

## MATERIALS AND METHODS

### Animals

Zebrafish (*Danio rerio* H.) were obtained from the Zebrafish International Resource Center and mated, staged, raised and processed as described (Westerfield, 2000). The lines *Tg(mpx:eGFP)^i114^* (Renshaw et al., 2006), *Tg(lyz:DsRED2)^nz50^ (Hall et al., 2007)*, *Tg(mfap4.1:mCherry-F)^xt12^* (Walton et al., 2015), *Tg(NFkB-RE:eGFP)^sh235^* (Kanther et al., 2011), *Tg(il1b:GFP-F)^ump3^* (Nguyen-Chi et al., 2014), *Tg(mfap4.1:mCherry-F), Tg(tnfa:eGFP-F)^ump5^* (Nguyen-Chi et al., 2015), *myd88*^hu3568/hu3568^ knockout fish (van der Vaart et al., 2012), and *mitfa^w2/w2^; mpv17^a9/a9^*, referred to here as casper, (White et al., 2008) were previously described. All the experiments complied with the Guidelines of the European Union Council (Directive 2010/63/EU) and Spanish Royal Decree, RD 53/2013. Experiments and procedures were performed as approved by the Bioethical Committees of the University of Murcia (approval numbers #395/2017).

### Silica crystal injection and gene edition

SiO_2_ crystals free of bacterial contamination (Nano-SiO_2_, diameter less than 100 nm, InvivoGen, #tlrl-sio-2) at a concentration of 10 mg/ml supplemented with phenol red, were injected into the hindbrain vesicle of 2 dpf larvae (Video S1).

The crRNA for zebrafish *nlrp3*, *gsdmea*/*gsdmeb* and *il1b* (Table S1), and tracrRNA were resuspended to reach 100 µM Nuclease-Free Duplex Buffer. One microliter of each was mixed and incubated for 5 min at 95 ^°^C for duplexing. When removed from the heat and cooled to room temperature, 1.43 µl of Nuclease-Free Duplex Buffer was added to the duplex giving a final concentration of 1000 ng/µl. Finally, the injection mix was prepared mixing 1 µl of duplex, 2.55 µl of Nuclease-Free Duplex Buffer, 0.25 µl Cas9 Nuclease V3 (IDT, #1081058) and 0.25 µl of phenol red, giving the final concentrations of 250 ng/µl of gRNA duplex and 500 ng/µl of Cas9. The prepared mix was microinjected into the yolk sac of one- to eight-cell-stage embryos using a microinjector (Narishige) (0.5–1 nl per embryo). The same amounts of gRNA were used in all experimental groups. The efficiency of gRNA was checked by analyzing the sequence of the target sequence with the ICE CRISPR Analysis Tool (Conant et al., 2022) after amplification using a specific pair of primers (Table S2).

### Survival and touch-evoke response assays

After injection of the SiO_2_ crystals into the hindbrain under anesthesia, the larvae were allowed to recover in egg water at 28-29 ^0^C and monitored for clinical signs of disease or mortality. The number of dead larvae was counted daily until 5^th^ day post injection (7^th^ dpf). At least three independent experiments were performed with a total number of 100-120 larvae per treatment.

The touch-evoke response assay was performed by delivering a mechanosensory stimulus to the embryo by touching it gently with a blunt needle on the top of the head (Sztal et al., 2016).

### Chemical treatments

In some experiments, 2 dpf larvae were manually dechorionated and treated for times reanging from 12 h to 5 d at 28 ^°^C by bath immersion with caspase-1 inhibitor VX-765 (100 µM), the CXCR2 inhibitor SB225002 (5 µM), the VEGFR1/2/3, FGFR1/2/3 and PDGFRα/β inhibitor nintedanib (10 nM-10 μM), the chemokine and collagen inhibitor pirfenidone (50 nM-500 μM), and/or the type-I collagen synthesis inhibitor halofuginone hydrobromide (10 nM-10 μM), all from Medical Chemical Express, diluted in egg water supplemented with 0.1% DMSO. In some experiments, 24 hpf embryos were treated with 0.3 % N-Phenylthiourea (PTU) in order to inhibit melanogenesis for *in vivo* imaging.

### Analysis of gene expression

Total RNA was extracted from the whole head/tail part of the zebrafish body with TRIzol reagent (Invitrogen) following the manufacturer’s instructions and treated with DNase I, amplification grade (1 U/mg RNA: Invitrogen). SuperScript IV RNase H Reverse Transcriptase (Invitrogen) was used to synthesize first-strand cDNA with random primer from 1 mg of total RNA at 50 ^°^C for 50 min. Real-time PCR was performed with an ABIPRISM 7500 instrument (Applied Biosystems) using SYBR Green PCR Core Reagents (Applied Biosystems). Reaction mixtures were incubated for 10 min at 95 ^°^C, followed by 40 cycles of 15 s at 95 ^°^C, 1 min at 60 ^°^C, and finally 15 s at 95 ^°^C, 1 min 60 ^°^C, and 15 s at 95 ^°^C. For each mRNA, gene expression was normalized to the ribosomal protein S11 gene (*rps11*) content in each sample using the Pfaffl method (Pfaffl, 2001). The primers used are shown in Table S2. In all cases, each PCR was performed with triplicate samples and repeated at least twice with independent samples.

### Caspase-1 activity assays

The caspase-1 activity was determined with the fluorometric substrate Z-YVAD 7-Amido-4-trifluoromethylcoumarin (Z-YVAD-AFC, caspase-1 substrate VI, Calbiochem), as described previously (Angosto et al., 2012; Lopez-Castejon et al., 2008), using 35–45 heads or the rest of the bodies of 3 dpf larvae. A representative caspase-1 activity graph from three biological replicates is shown in the figures.

### *In vivo* imaging

To study immune cell recruitment to the injection site and the Nfkb, Il1b and Tnfα activation pattern, 2 dpf *mpx:eGFP*, *mfap4:mcherry*, *nfkb:eGFP*, *il1b:eGFP* and *tnfa:eGFP* zebrafish larvae were anaesthetized in embryo medium with 0.16 mg/ml buffered tricaine. SiO_2_ crystal-injected larvae were imaged (both hindbrain and the whole-body area) at 1, 3, 6, 9, 24 and 48 hours hpi using a Leica MZ16F fluorescence stereo microscope. The number of neutrophils or macrophages was visually countd in blinded samples, and the fluorescence intensity was obtained and analyzed with ImageJ (FIJI) software (Schindelin et al., 2012).

Macrophage polarization was evaluated in *Tg(mpeg1:mCherry, tnfa:eGFP)* (Nguyen-Chi et al., 2015). Briefly, 2 dpf larvae were anaesthetized in embryo medium with 0.16 mg/ml buffered tricaine. Nano-SiO_2_ at a concentration of 10 mg/ml supplemented with phenol red, were injected into the hindbrain vesicle. Images of the head and the whole-body area were taken at 6 hpi using a Leica MZ16F fluorescence stereo microscope. The number of macrophages in the whole body or in the head (mCherry^+^) and *tnfa*-expressing macrophages (eGFP^+^) was visually counted in blinded samples. The ratio between Tnfa^+^ and Tnfa^-^ macrophages was calculated.

### Statistical analysis

Data are shown as mean ± s.e.m. and were analyzed by one-way analysis of variance and a Tukey multiple range test to determine differences between groups. The differences between two samples were analyzed by Student’s *t*-test. A log-rank test was used to calculate statistical differences in the survival of the different experimental groups.

## ACKNOWLEDGMENTS

This work has been funded by Fundación Séneca, CARM, Spain (research grant 20793/PI/18 to VM and Saavedra Fajardo postdoctoral contract to SC), the European Union Horizon 2020 research and innovation program under the Marie Skłodowska-Curie Actions (grant agreement No.955576 – INFLANET), the Spanish Ministry of Science and Innovation (Juan de la Cierva-Incorporación postdoctoral contract to SDT), co-funded with European Regional Development Funds. We thank I. Fuentes and P. Martinez for their excellent technical assistance, and Profs Tobin, Crosier, Renshaw, Zon, Meijer and Lutfalla for the zebrafish lines.

## AUTHOR CONTRIBUTIONS

The authors offer the following declarations about their contributions: Conceived and designed the experiments: SDT, VM. Performed the experiments: SDT, AP, AM-L. Analyzed the data: SDT, AP, AM-L JAR-L, PMdC, SC, VM. Wrote the paper: SDT, VM.

---

**Figure S1 (related to all figures):**
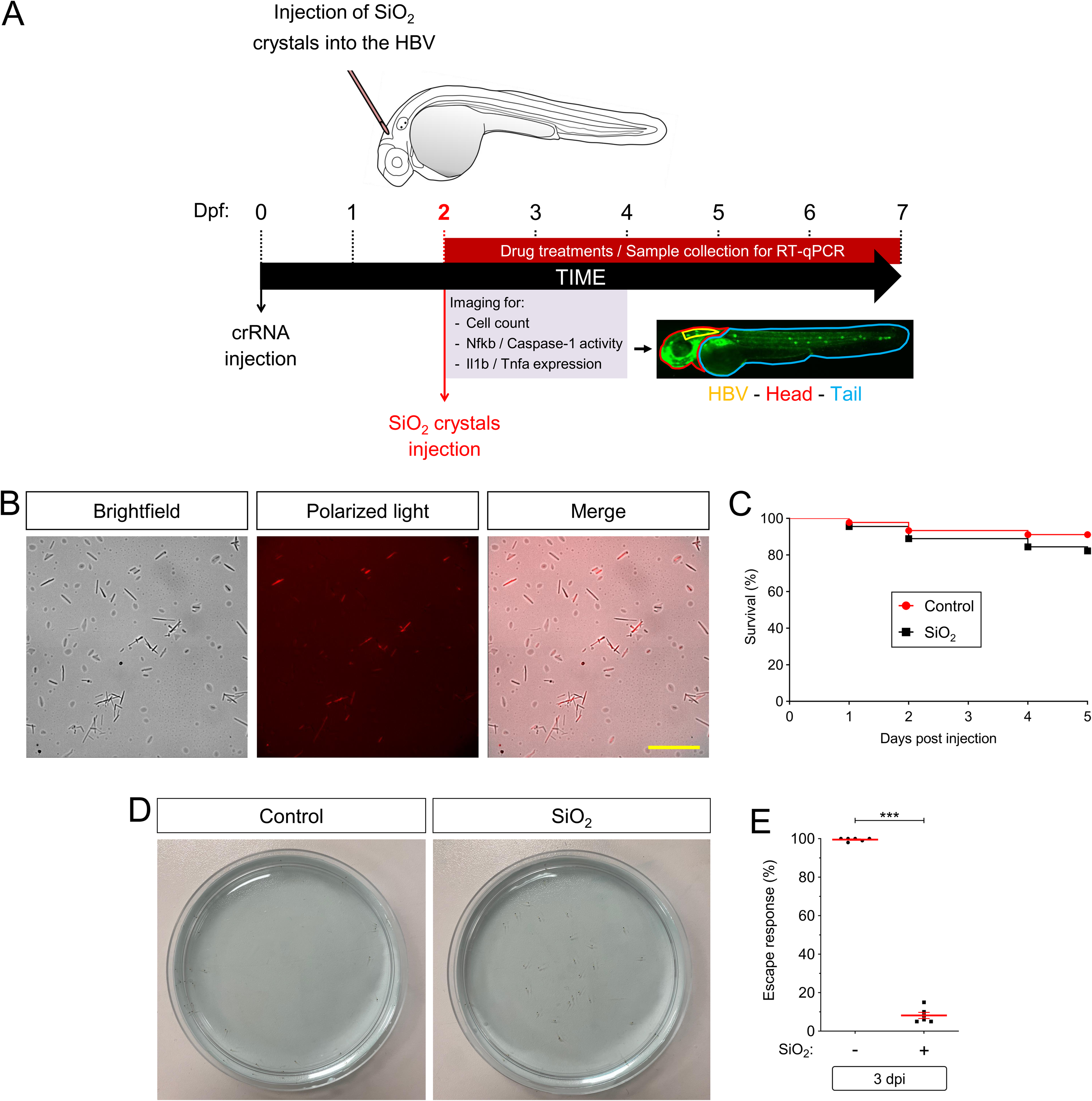
Experimental design and effect of silica crystals in zebrafish larval survival and behavior. (A) Scheme showing the experimental design for all experiments, including SiO_2_ injection, gene editing and chemical inhibition. (B) Micrographs of SiO_2_ crystals observed under brightfield and polarized light. (C) Larval survival after SiO_2_ injection. (D, E) Touch-evoke response of 3 dpf larvae injected with PBS or SiO_2_. Each dot represents one individual and the mean ± S.E.M. for each group is also shown. P values were calculated using log-rank test (C) and Student t test (E). ***p≤0.001. Bar: 50 µm.

**Figure S2 (related to Figure 3).**
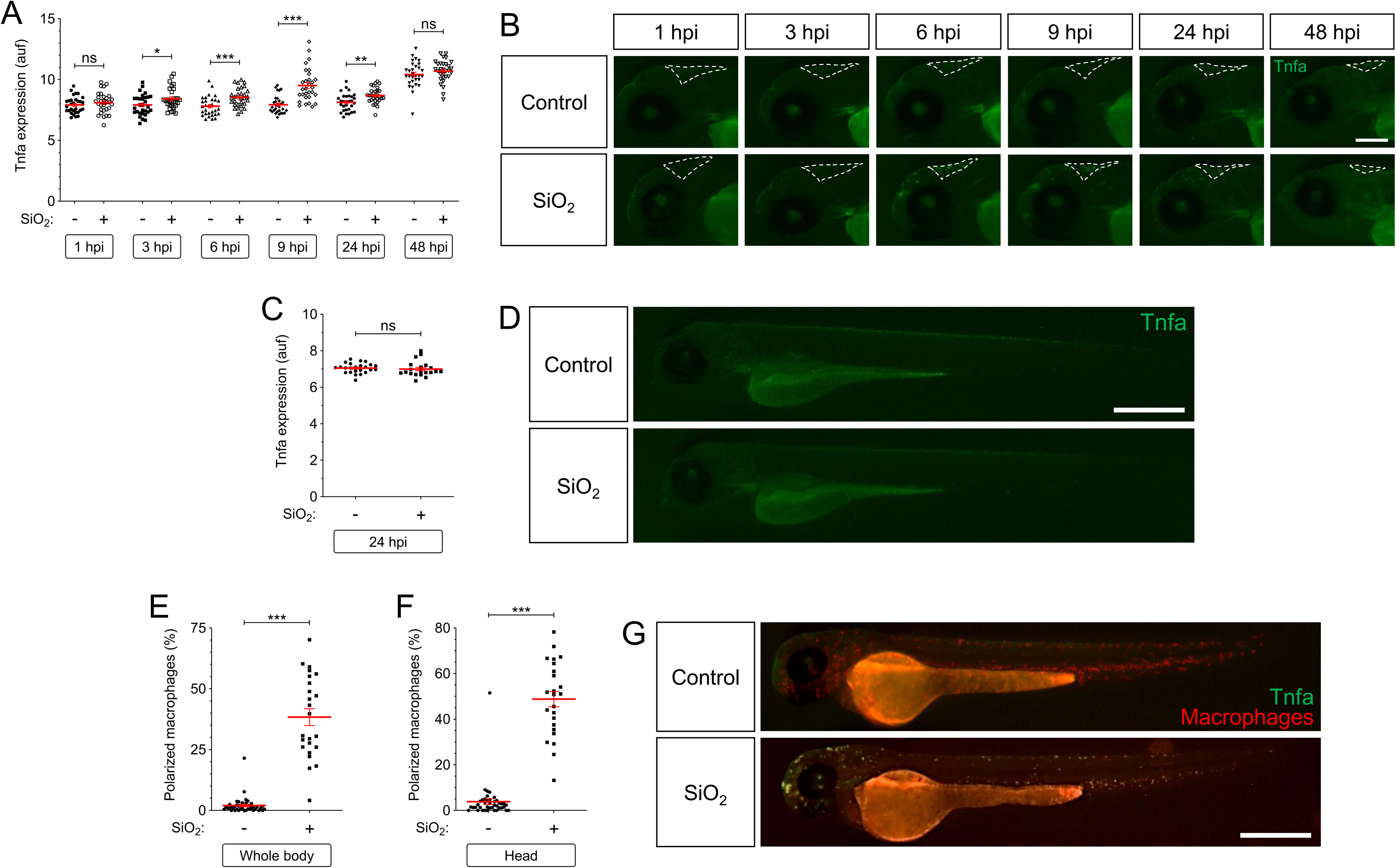
Silica crystals promote M1 polarization of macrophage in zebrafish. SiO_2_ crystals or PBS were injected in the hindbrain ventricle (HBV, are indicated with a dashed line) of 2 dpf *Tg(mpeg1:mCherry-F); Tg(tnfa:eGFP)*. Tnfa expression (A-D) and M1 macrophages (Cherry^+^/Tnfa^+^) were analyzed at the indicated times by fluorescence microscopy. Each dot represents one individual and the mean ± S.E.M. for each group is also shown. P values were calculated using one-way ANOVA and Tukey multiple range test. ns, not significant, *≤p0.05, **p≤0.01, ***p≤0.001. Bar: 100 µm (B), 500 µm (D, G).

**Figure S3 (related to Figure 3).**
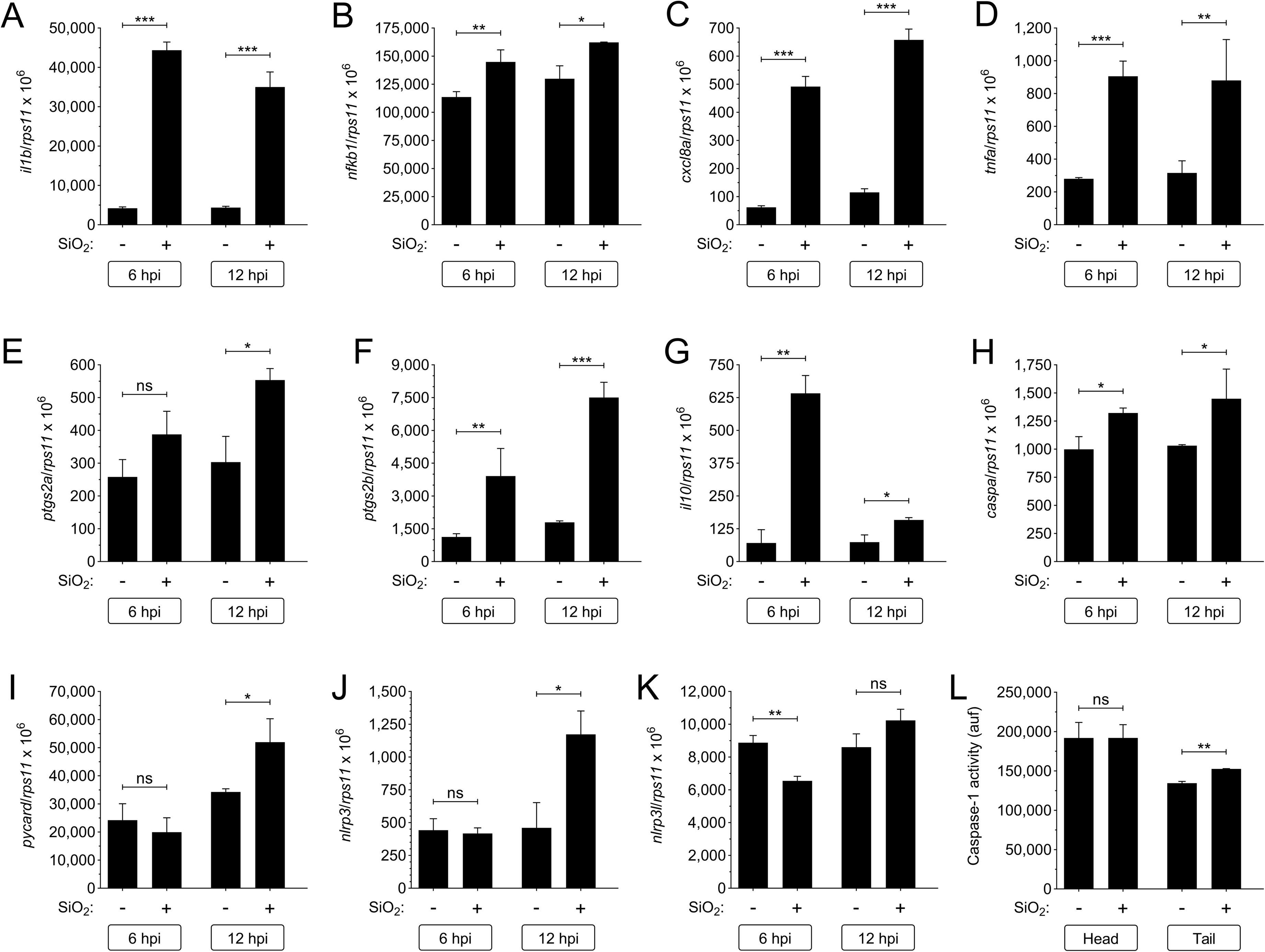
Silica crystals induce expression of genes encoding inflammatory mediators in zebrafish. SiO_2_ crystals or PBS were injected in the hindbrain ventricle (HBV) of 2 dpf wild type larvae. The transcript levels of the indicated genes were analyzed at 6 and 12 hpi by RT-qPCR in larval head (A-K), while caspase-1 activity was determined at 24 hpi using a fluorogenic substrate (L). The data are shown as the mean ± S.E.M. P values were calculated using one-way ANOVA and Tukey multiple range test. ns, not significant, *≤p0.05, **p≤0.01, ***p≤0.001. auf, arbitrary units of fluorescence.

**Figure S4 (related to Figure 6).**
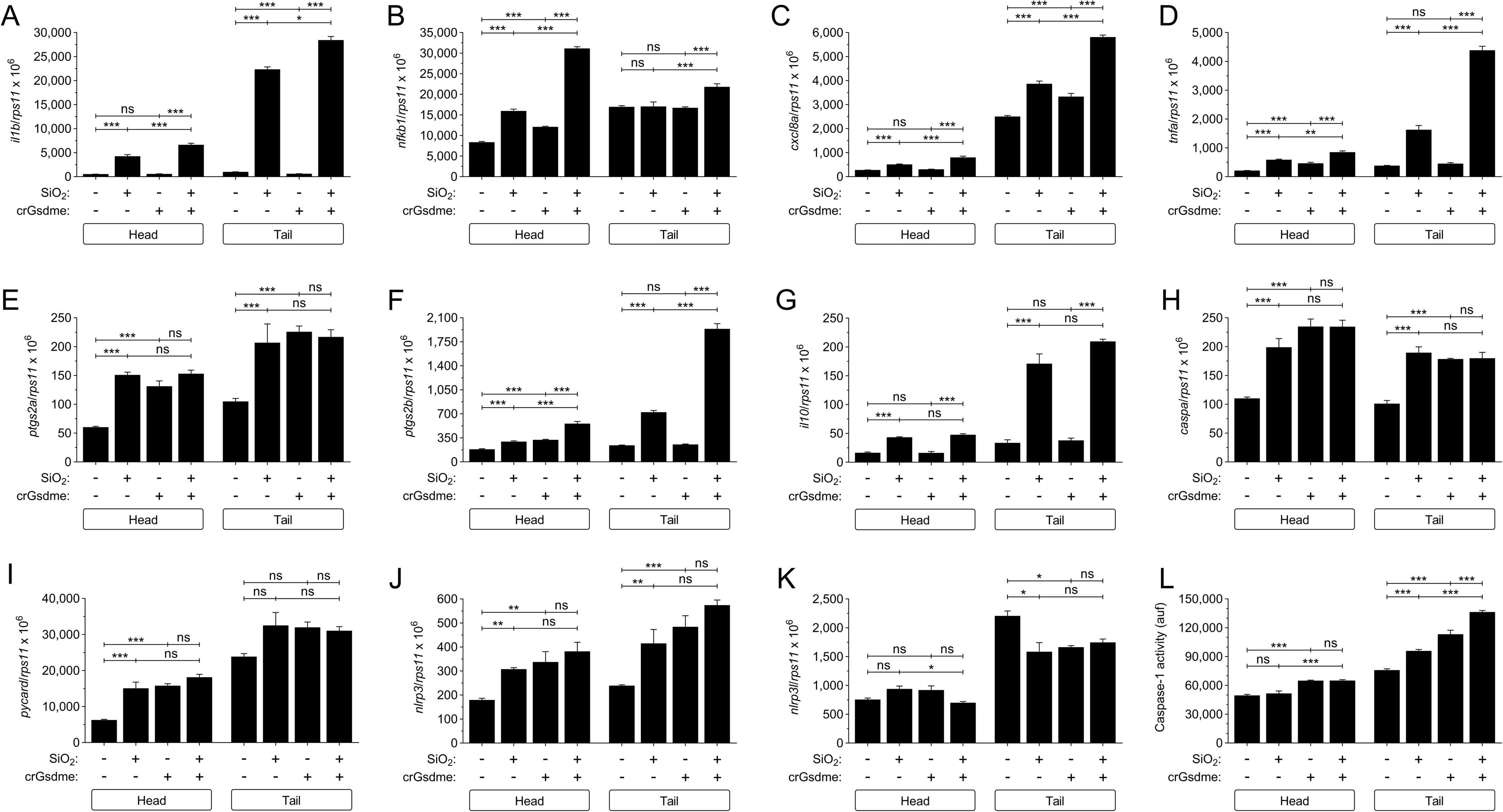
Gsdmea/b deficiency enhances silica crystal-induced inflammation in zebrafish. Wild type 1-8 cell stage embryos were microinjected with control or *gsdmea/b* crRNA/Cas9 complexes. At 2dpf, SiO_2_ crystals or PBS were injected in the hindbrain ventricle (HBV). The transcript levels of the indicated genes were analyzed at 12 hpi by RT-qPCR in larval head (A-K), while caspase-1 activity was determined at 24 hpi using a fluorogenic substrate (L). The data are shown as the mean ± S.E.M. P values were calculated using one-way ANOVA and Tukey multiple range test. ns, not significant, *≤p0.05, **p≤0.01, ***p≤0.001. auf, arbitrary units of fluorescence.

**Figure S5 (related to Figure 6).**
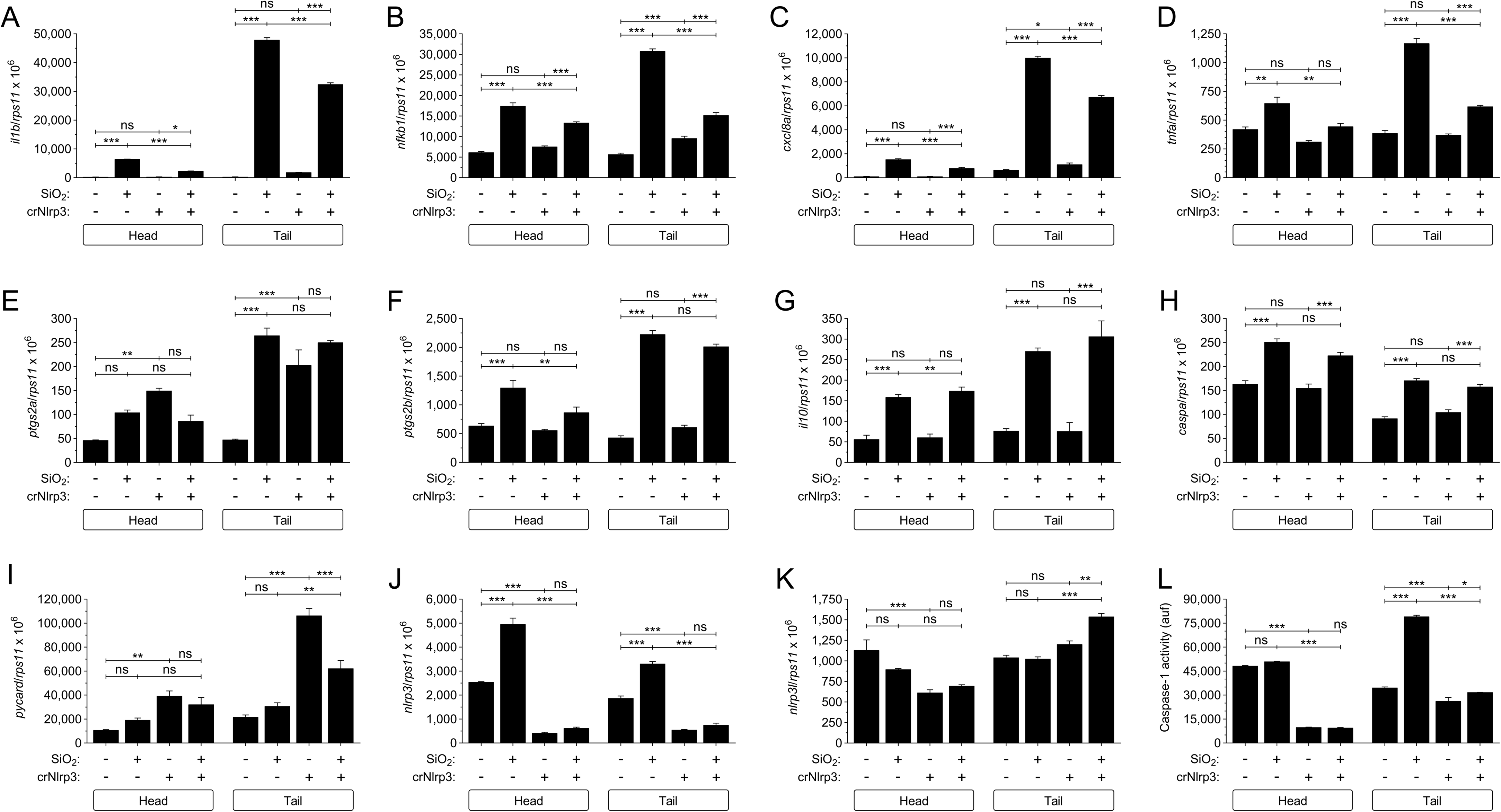
Nlrp3 deficiency attenuates silica crystal-induced inflammation in zebrafish. Wild type 1-8 cell stage embryos were microinjected with control or *nlrp3* crRNA/Cas9 complexes. At 2dpf, SiO_2_ crystals or PBS were injected in the hindbrain ventricle (HBV). The transcript levels of the indicated genes were analyzed at 12 hpi by RT-qPCR in larval head (A-K), while caspase-1 activity was determined at 24 hpi using a fluorogenic substrate (L). The data are shown as the mean ± S.E.M. P values were calculated using one-way ANOVA and Tukey multiple range test. ns, not significant, *≤p0.05, **p≤0.01, ***p≤0.001. auf, arbitrary units of fluorescence.

**Figure S6 (related to Figure 6).**
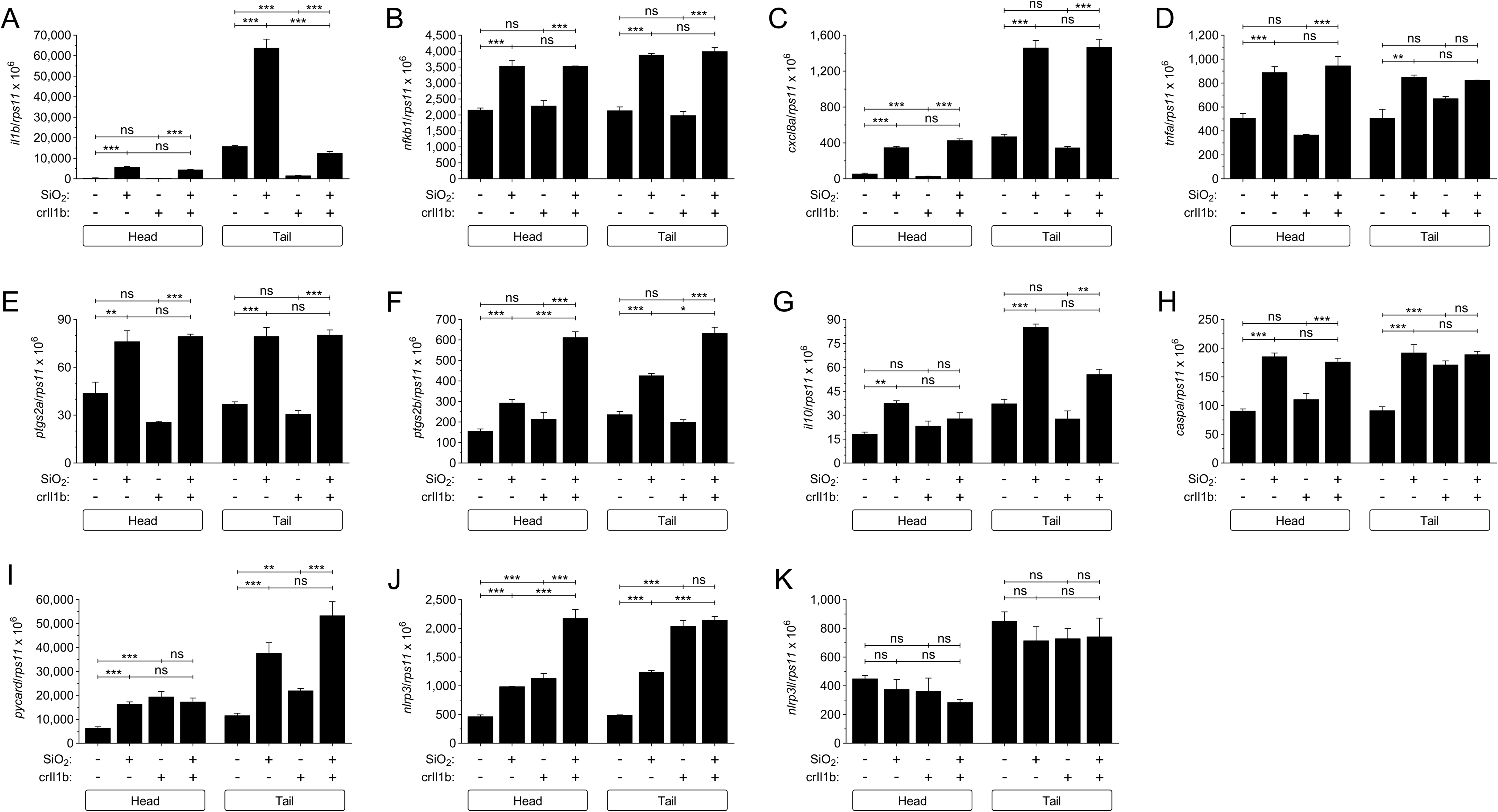
Il1b deficiency hardly affects silica crystal-induced inflammation in zebrafish. Wild type 1-8 cell stage embryos were microinjected with control or *il1b* crRNA/Cas9 complexes. At 2dpf, SiO_2_ crystals or PBS were injected in the hindbrain ventricle (HBV). The transcript levels of the indicated genes were analyzed at 12 hpi by RT-qPCR in larval head (A-K). The data are shown as the mean ± S.E.M. P values were calculated using one-way ANOVA and Tukey multiple range test. ns, not significant, *≤p0.05, **p≤0.01, ***p≤0.001.

**Figure S7 (related to Figure 8):**
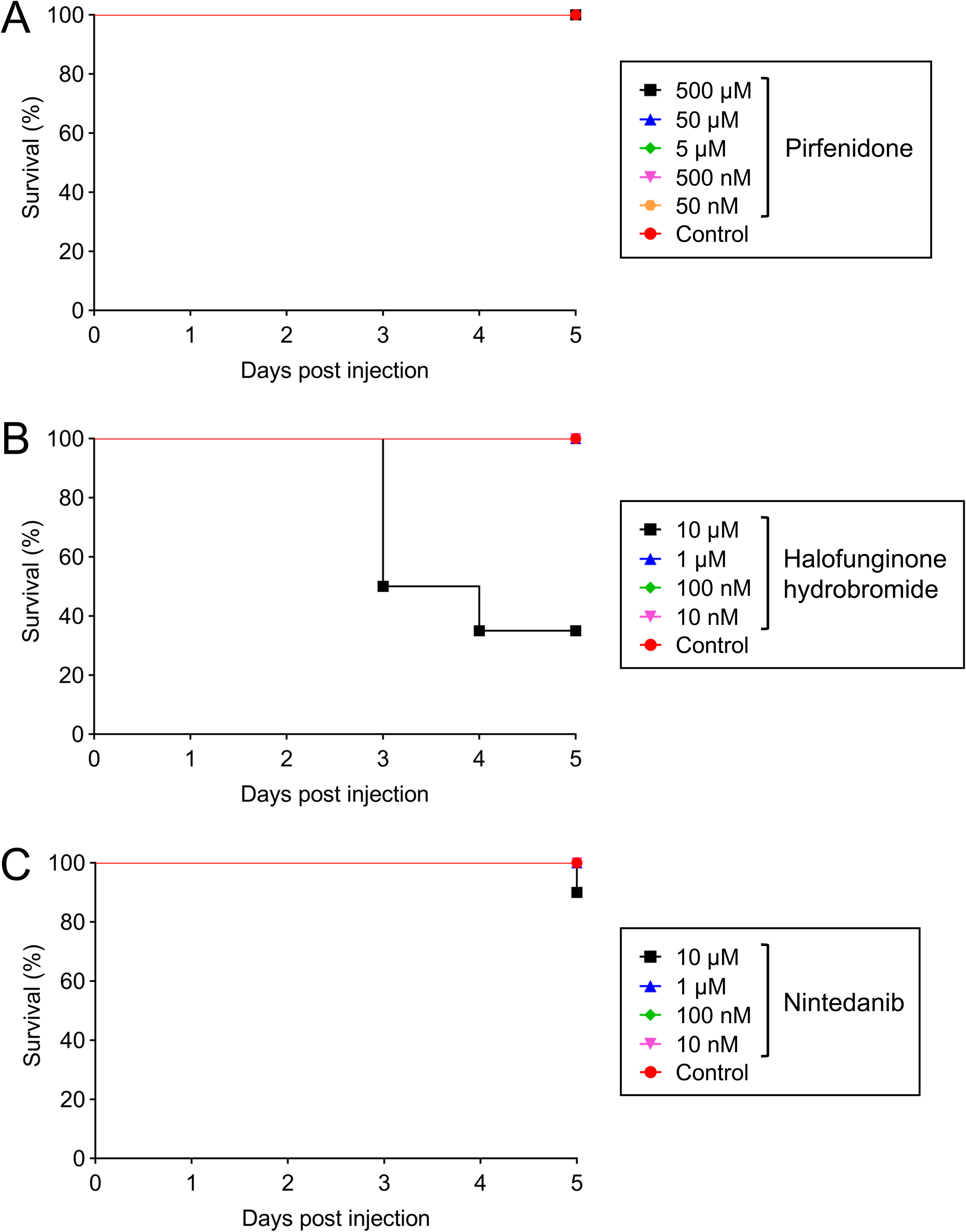
Survival of zebrafish larvae treated with different concentrations of antifibrotic drugs. Kaplan-Meier survival curves of wild type larvae treated by bath immersion with the indicated concentrations of the antifibrotic drugs pirfenidone (A), nintedanib (B) or halofuginone hydrobromide (C).

**Figure S8 (related to Figure 8):**
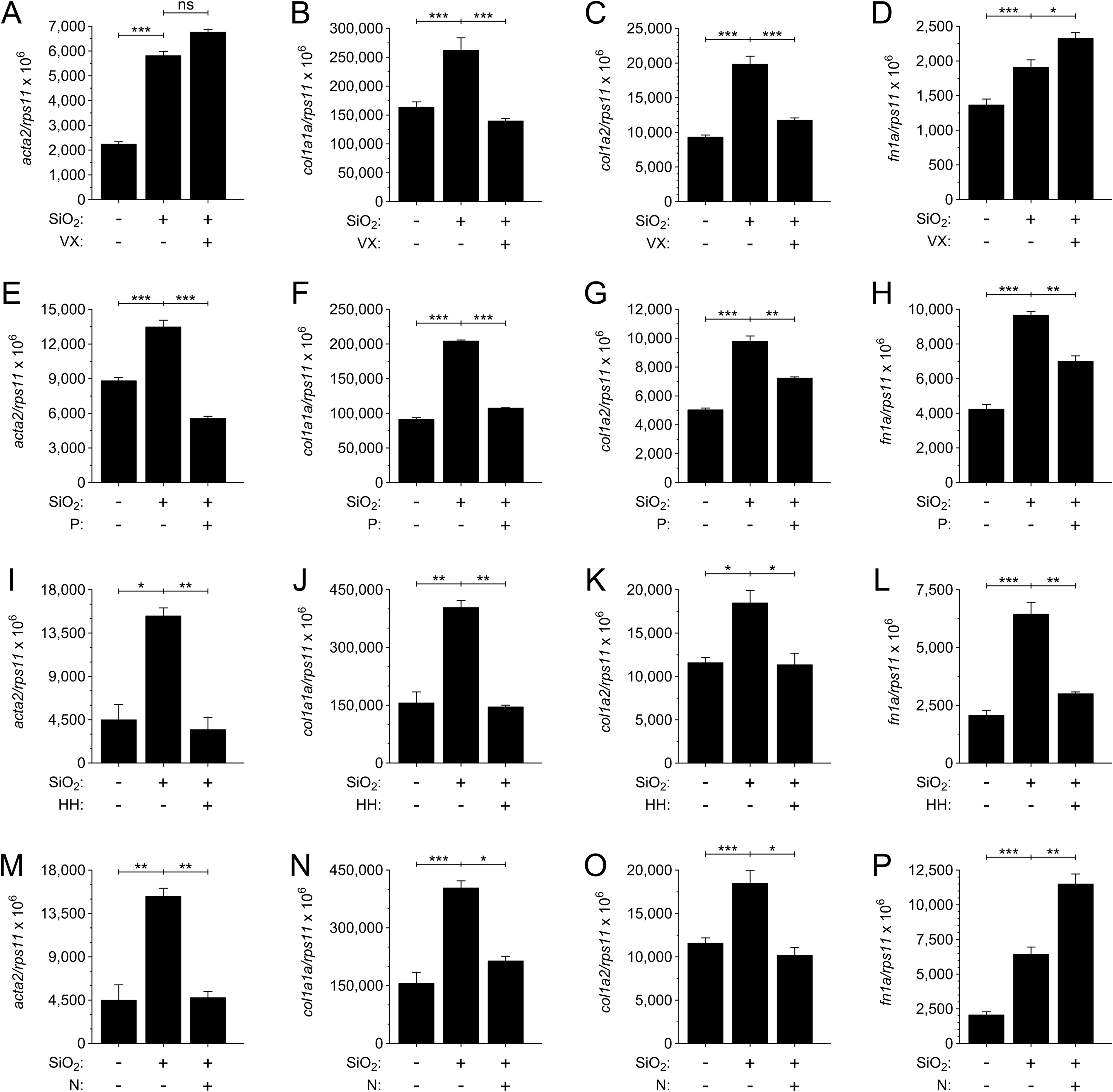
Silica crystals promote fibrosis in zebrafish. SiO_2_ crystals or PBS were injected in the hindbrain ventricle (HBV) of 2 dpf wild type larvae in the presence of DMSO, the casapase-1 inhibitor VX-765 (VX), or the antifibrotic drugs pirfenidone (P), halofuginone hydrobromide (HH) or nintedanib (N). The transcript levels of the indicated genes encoding fibrosis biomarkers were analyzed at 120 hpi by RT-qPCR in larval head. The data are shown as the mean ± S.E.M. P values were calculated using one-way ANOVA and Tukey multiple range test. ns, not significant, *≤p0.05, **p≤0.01, ***p≤0.001.

**Video S1*: Experimental procedure used to microinject silica crystals into the hindbrain ventricle of 2 dpf zebrafish larvae*.**

**Table S1.**
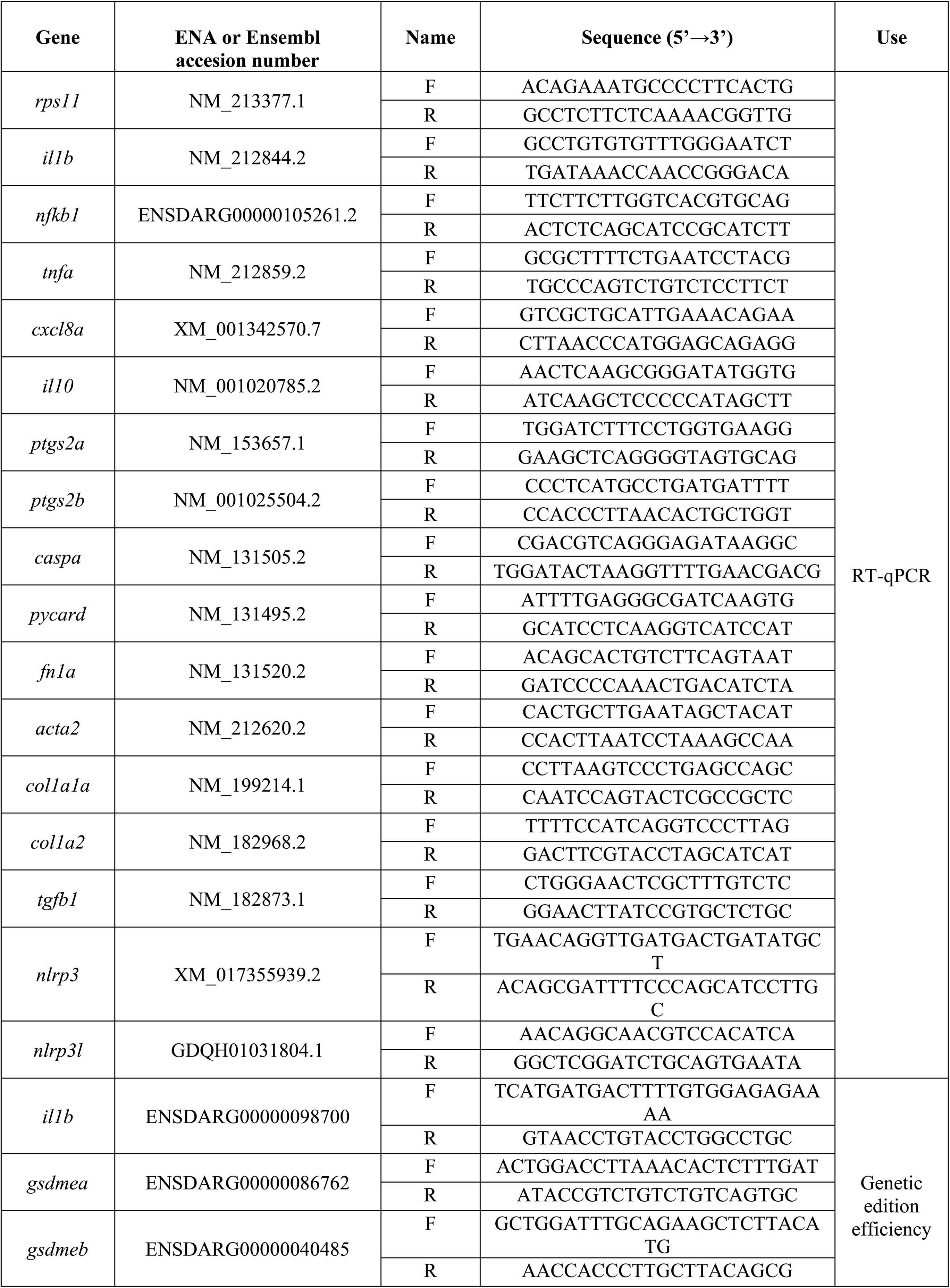
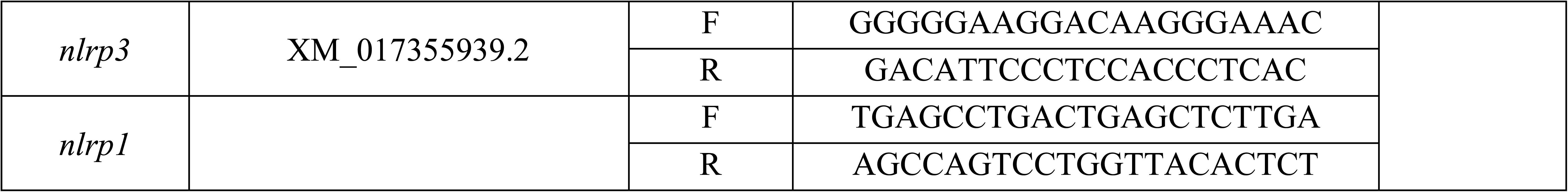
Primers used in this study. The gene symbols followed the Zebrafish Nomenclature Guidelines (http://zfin.org/zf_info/nomen.html). ENA, European Nucleotide Archive.

**Table S2.**
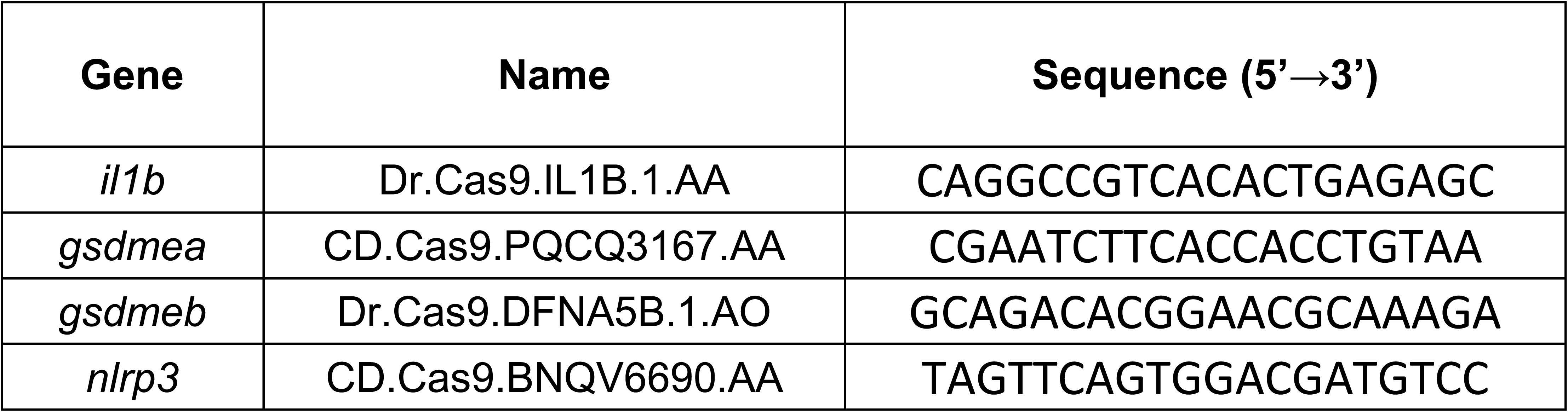
crRNA used in this study (IDT). The gene symbols followed the Zebrafish Nomenclature Guidelines (http://zfin.org/zf_info/nomen.html).

## REFERENCES

Allison, S.J. 2014. Fibrosis: Regulation of fibrotic signalling by TGF-β receptor tyrosine phosphorylation. Nat Rev Nephrol 10:484.

Angosto, D., G. López-Castejón, A. López-Muñoz, M.P. Sepulcre, M. Arizcun, J. Meseguer, and V. Mulero. 2012. Evolution of inflammasome functions in vertebrates: Inflammasome and caspase-1 trigger fish macrophage cell death but are dispensable for the processing of IL-1β. Innate Immun 18:815–824.

Barnes, H., N.S.L. Goh, T.L. Leong, and R. Hoy. 2019. Silica-associated lung disease: An old-world exposure in modern industries. Respirology 24:1165–1175.

Beamer, C.A., and A. Holian. 2008. Silica suppresses Toll-like receptor ligand-induced dendritic cell activation. FASEB J 22:2053–2063.

Bournele, D., and D. Beis. 2016. Zebrafish models of cardiovascular disease. Heart Fail Rev 21:803–813.

Bowden, D.H., C. Hedgecock, and I.Y. Adamson. 1989. Silica-induced pulmonary fibrosis involves the reaction of particles with interstitial rather than alveolar macrophages. J Pathol 158:73–80.

Broz, P., P. Pelegrín, and F. Shao. 2020. The gasdermins, a protein family executing cell death and inflammation. Nat Rev Immunol 20:143–157.

Cambier, C.J., K.K. Takaki, R.P. Larson, R.E. Hernandez, D.M. Tobin, K.B. Urdahl, C.L. Cosma, and L. Ramakrishnan. 2014. Mycobacteria manipulate macrophage recruitment through coordinated use of membrane lipids. Nature 505:218–222.

Cao, Z., M. Song, Y. Liu, J. Pang, Z. Li, X. Qi, T. Shu, B. Li, D. Wei, J. Chen, J. Wang, and C. Wang. 2020. A novel pathophysiological classification of silicosis models provides some new insights into the progression of the disease. Ecotoxicol Environ Saf 202:110834.

Cao, Z.J., Y. Liu, Z. Zhang, P.R. Yang, Z.G. Li, M.Y. Song, X.M. Qi, Z.F. Han, J.L. Pang, B.C. Li, X.R. Zhang, H.P. Dai, J. Wang, and C. Wang. 2022. Pirfenidone ameliorates silica-induced lung inflammation and fibrosis in mice by inhibiting the secretion of interleukin-17A. Acta Pharmacol Sin 43:908–918.

Carrieri, M., C. Guzzardo, D. Farcas, and L.G. Cena. 2020. Characterization of Silica Exposure during Manufacturing of Artificial Stone Countertops. Int J Environ Res Public Health 17:

Cassel, S.L., S.C. Eisenbarth, S.S. Iyer, J.J. Sadler, O.R. Colegio, L.A. Tephly, A.B. Carter, P.B. Rothman, R.A. Flavell, and F.S. Sutterwala. 2008. The Nalp3 inflammasome is essential for the development of silicosis. Proc Natl Acad Sci U S A 105:9035–9040.

Cavalli, G., F. Fallanca, C.A. Dinarello, and L. Dagna. 2015. Treating pulmonary silicosis by blocking interleukin 1. Am J Respir Crit Care Med 191:596–598.

Chan, J.Y.W., J.C.C. Tsui, P.T.W. Law, W.K.W. So, D.Y.P. Leung, M.M.K. Sham, S.K.W. Tsui, and C.W.H. Chan. 2018. Regulation of TLR4 in silica-induced inflammation: An underlying mechanism of silicosis. Int J Med Sci 15:986–991.

Chen, H., X. Wu, Z. Gu, S. Chen, X. Zhou, Y. Zhang, Q. Liu, Z. Wang, and D. Yang. 2021. Zebrafish gasdermin E cleavage-engaged pyroptosis by inflammatory and apoptotic caspases. Dev Comp Immunol 124:104203.

Chen, S., B. Han, X. Geng, P. Li, M.F. Lavin, A.J. Yeo, C. Li, J. Sun, C. Peng, H. Shao, and Z. Du. 2022. Microcrystalline silica particles induce inflammatory response via pyroptosis in primary human respiratory epithelial cells. Environ Toxicol 37:385–400.

Conant, D., T. Hsiau, N. Rossi, J. Oki, T. Maures, K. Waite, J. Yang, S. Joshi, R. Kelso, K. Holden, B.L. Enzmann, and R. Stoner. 2022. Inference of CRISPR Edits from Sanger Trace Data. CRISPR J 5:123–130.

Davis, G.S., L.M. Pfeiffer, K.E. Leslie, and D.R. Hemenway. 1996. Macrophage-lymphocyte cytokine interactions in silicosis. Chest 109:49S–50S.

de Oliveira, S., A. Lopez-Munoz, F.J. Martinez-Navarro, J. Galindo-Villegas, V. Mulero, and A. Calado. 2015. Cxcl8-l1 and Cxcl8-l2 are required in the zebrafish defense against Salmonella Typhimurium. Dev Comp Immunol 49:44–48.

Diniz, M.S., R. Salgado, V.J. Pereira, G. Carvalho, A. Oehmen, M.A. Reis, and J.P. Noronha. 2015. Ecotoxicity of ketoprofen, diclofenac, atenolol and their photolysis byproducts in zebrafish (Danio rerio). Sci Total Environ 505:282–289.

Dostert, C., V. Pétrilli, R. Van Bruggen, C. Steele, B.T. Mossman, and J. Tschopp. 2008. Innate immune activation through Nalp3 inflammasome sensing of asbestos and silica. Science 320:674–677.

Duchange, L., A. Brichet, C. Lamblin, I. Tillie, A.B. Tonnel, and B. Wallaert. 1998. [Acute silicosis. Clinical, radiologic, functional, and cytologic characteristics of the broncho-alveolar fluids. Observations of 6 cases]. Rev Mal Respir 15:527–534.

Edwards, G. 2019. Accelerated Silicosis-An Emerging Epidemic Associated with Engineered Stone. Comment on Leso, V. et al. Artificial Stone Associated Silicosis: A Systematic Review. Int J Environ Res Public Health 16:

Espín-Palazón, R., A. Martínez-López, F.J. Roca, A. López-Muñoz, S.D. Tyrkalska, S. Candel, D. García-Moreno, A. Falco, J. Meseguer, A. Estepa, and V. Mulero. 2016. TNFα Impairs Rhabdoviral Clearance by Inhibiting the Host Autophagic Antiviral Response. PLoS Pathog 12:e1005699.

Ferreira, T.P., L.L. Mariano, R. Ghilosso-Bortolini, A.C. de Arantes, A.J. Fernandes, M. Berni, V. Cecchinato, M. Uguccioni, R. Maj, A. Barberis, P.M. Silva, and M.A. Martins. 2016. Potential of PEGylated Toll-Like Receptor 7 Ligands for Controlling Inflammation and Functional Changes in Mouse Models of Asthma and Silicosis. Front Immunol 7:95.

Gabellini, C., E. Gomez-Abenza, S. Ibanez-Molero, M.G. Tupone, A.B. Perez-Oliva, S. de Oliveira, D. Del Bufalo, and V. Mulero. 2018. Interleukin 8 mediates bcl-xL-induced enhancement of human melanoma cell dissemination and angiogenesis in a zebrafish xenograft model. Int J Cancer 142:584–596.

Guo, J., Z. Yang, Q. Jia, C. Bo, H. Shao, and Z. Zhang. 2019. Pirfenidone inhibits epithelial-mesenchymal transition and pulmonary fibrosis in the rat silicosis model. Toxicol Lett 300:59–66.

Ha, H., B. Debnath, and N. Neamati. 2017. Role of the CXCL8-CXCR1/2 Axis in Cancer and Inflammatory Diseases. Theranostics 7:1543–1588.

Hall, C., M.V. Flores, T. Storm, K. Crosier, and P. Crosier. 2007. The zebrafish lysozyme C promoter drives myeloid-specific expression in transgenic fish. BMC Dev Biol 7:42.

Hamilton, R.F., S.A. Thakur, and A. Holian. 2008. Silica binding and toxicity in alveolar macrophages. Free Radic Biol Med 44:1246–1258.

Harijith, A., D.L. Ebenezer, and V. Natarajan. 2014. Reactive oxygen species at the crossroads of inflammasome and inflammation. Front Physiol 5:352.

Hason, M., and P. Bartůněk. 2019. Zebrafish Models of Cancer-New Insights on Modeling Human Cancer in a Non-Mammalian Vertebrate. Genes (Basel) 10:

Hegde, B., S.R. Bodduluri, S.R. Satpathy, R.S. Alghsham, V.R. Jala, S.M. Uriarte, D.H. Chung, M.B. Lawrenz, and B. Haribabu. 2018. Inflammasome-Independent Leukotriene B4 Production Drives Crystalline Silica-Induced Sterile Inflammation. J Immunol 200:3556–3567.

Herbomel, P., B. Thisse, and C. Thisse. 1999. Ontogeny and behaviour of early macrophages in the zebrafish embryo. Development 126:3735–3745.

Honnons, S., and J.M. Porcher. 2000. In vivo experimental model for silicosis. J Environ Pathol Toxicol Oncol 19:391–400.

Hornung, V., F. Bauernfeind, A. Halle, E.O. Samstad, H. Kono, K.L. Rock, K.A. Fitzgerald, and E. Latz. 2008. Silica crystals and aluminum salts activate the NALP3 inflammasome through phagosomal destabilization. Nat Immunol 9:847–856.

Howe, K., M.D. Clark, C.F. Torroja, J. Torrance, C. Berthelot, M. Muffato, J.E. Collins, S. Humphray, K. McLaren, L. Matthews, S. McLaren, I. Sealy, M. Caccamo, C. Churcher, C. Scott, J.C. Barrett, R. Koch, G.J. Rauch, S. White, W. Chow, B. Kilian, L.T. Quintais, J.A. Guerra-Assunção, Y. Zhou, Y. Gu, J. Yen, J.H. Vogel, T. Eyre, S. Redmond, R. Banerjee, J. Chi, B. Fu, E. Langley, S.F. Maguire, G.K. Laird, D. Lloyd, E. Kenyon, S. Donaldson, H. Sehra, J. Almeida-King, J. Loveland, S. Trevanion, M. Jones, M. Quail, D. Willey, A. Hunt, J. Burton, S. Sims, K. McLay, B. Plumb, J. Davis, C. Clee, K. Oliver, R. Clark, C. Riddle, D. Elliot, D. Eliott, G. Threadgold, G. Harden, D. Ware, S. Begum, B. Mortimore, B. Mortimer, G. Kerry, P. Heath, B. Phillimore, A. Tracey, N. Corby, M. Dunn, C. Johnson, J. Wood, S. Clark, S. Pelan, G. Griffiths, M. Smith, R. Glithero, P. Howden, N. Barker, C. Lloyd, C. Stevens, J. Harley, K. Holt, G. Panagiotidis, J. Lovell, H. Beasley, C. Henderson, D. Gordon, K. Auger, D. Wright, J. Collins, C. Raisen, L. Dyer, K. Leung, L. Robertson, K. Ambridge, D. Leongamornlert, S. McGuire, R. Gilderthorp, C. Griffiths, D. Manthravadi, S. Nichol, G. Barker, S. Whitehead, M. Kay, J. Brown, C. Murnane, E. Gray, M. Humphries, N. Sycamore, D. Barker, D. Saunders, J. Wallis, A. Babbage, S. Hammond, M. Mashreghi-Mohammadi, L. Barr, S. Martin, P. Wray, A. Ellington, N. Matthews, M. Ellwood, R. Woodmansey, G. Clark, J. Cooper, A. Tromans, D. Grafham, C. Skuce, R. Pandian, R. Andrews, E. Harrison, A. Kimberley, J. Garnett, N. Fosker, R. Hall, P. Garner, D. Kelly, C. Bird, S. Palmer, I. Gehring, A. Berger, C.M. Dooley, Z. Ersan-Ürün, C. Eser, H. Geiger, M. Geisler, L. Karotki, A. Kirn, J. Konantz, M. Konantz, M. Oberländer, S. Rudolph-Geiger, M. Teucke, C. Lanz, G. Raddatz, K. Osoegawa, B. Zhu, A. Rapp, S. Widaa, C. Langford, F. Yang, S.C. Schuster, N.P. Carter, J. Harrow, Z. Ning, J. Herrero, S.M. Searle, A. Enright, R. Geisler, R.H. Plasterk, C. Lee, M. Westerfield, P.J. de Jong, L.I. Zon, J.H. Postlethwait, C. Nüsslein-Volhard, T.J. Hubbard, H. Roest Crollius, J. Rogers, and D.L. Stemple. 2013. The zebrafish reference genome sequence and its relationship to the human genome. Nature 496:498–503.

Kanther, M., X. Sun, M. Mühlbauer, L.C. Mackey, E.J. Flynn, M. Bagnat, C. Jobin, and J.F. Rawls. 2011. Microbial colonization induces dynamic temporal and spatial patterns of NF-κB activation in the zebrafish digestive tract. Gastroenterology 141:197–207.

Kimmel, C.B., W.W. Ballard, S.R. Kimmel, B. Ullmann, and T.F. Schilling. 1995. Stages of embryonic development of the zebrafish. Dev Dyn 203:253–310.

Kramer, M.R., P.D. Blanc, E. Fireman, A. Amital, A. Guber, N.A. Rhahman, and D. Shitrit. 2012. Artificial stone silicosis [corrected]: disease resurgence among artificial stone workers. Chest 142:419–424.

Lapp, N.L., and V. Castranova. 1993. How silicosis and coal workers’ pneumoconiosis develop--a cellular assessment. Occup Med 8:35–56.

Leung, C.C., I.T. Yu, and W. Chen. 2012. Silicosis. Lancet 379:2008–2018.

Li, X.X., D.Y. Jiang, X.X. Huang, S.L. Guo, W. Yuan, and H.P. Dai. 2015. Toll-like receptor 4 promotes fibrosis in bleomycin-induced lung injury in mice. Genet Mol Res 14:17391–17398.

Lopes-Pacheco, M., E. Bandeira, and M.M. Morales. 2016. Cell-Based Therapy for Silicosis. Stem Cells Int 2016:5091838.

Lopez-Castejon, G., M.P. Sepulcre, I. Mulero, P. Pelegrin, J. Meseguer, and V. Mulero. 2008. Molecular and functional characterization of gilthead seabream Sparus aurata caspase-1: the first identification of an inflammatory caspase in fish. Mol Immunol 45:49–57.

Lozano-Gil, J.M., L. Rodriguez-Ruiz, S.D. Tyrkalska, D. Garcia-Moreno, A.B. Perez-Oliva, and V. Mulero. 2022. Gasdermin E mediates pyroptotic cell death of neutrophils and macrophages in a zebrafish model of chronic skin inflammation. Dev Comp Immunol 104404.

MacRae, C.A., and R.T. Peterson. 2015. Zebrafish as tools for drug discovery. Nat Rev Drug Discov 14:721–731.

Martin, K.R. 2007. The chemistry of silica and its potential health benefits. J Nutr Health Aging 11:94–97.

Mayeux, J.M., D.H. Kono, and K.M. Pollard. 2019. Development of experimental silicosis in inbred and outbred mice depends on instillation volume. Sci Rep 9:14190.

McElroy, A.N., R. Invernizzi, J.W. Laskowska, A. O’Neill, M. Doroudian, M. Moghoofei, S. Mostafaei, F. Li, A.A. Przybylski, D.N. O’Dwyer, A.G. Bowie, P.G. Fallon, T.M. Maher, C.M. Hogaboam, P.L. Molyneaux, N. Hirani, M.E. Armstrong, and S.C. Donnelly. 2022. Candidate Role for Toll-like Receptor 3 L412F Polymorphism and Infection in Acute Exacerbation of Idiopathic Pulmonary Fibrosis. Am J Respir Crit Care Med 205:550–562.

Mossman, B.T., and A. Churg. 1998. Mechanisms in the pathogenesis of asbestosis and silicosis. Am J Respir Crit Care Med 157:1666–1680.

Nemmar, A., B. Nemery, P.H. Hoet, N. Van Rooijen, and M.F. Hoylaerts. 2005. Silica particles enhance peripheral thrombosis: key role of lung macrophage-neutrophil cross-talk. Am J Respir Crit Care Med 171:872–879.

Newton, K., V.M. Dixit, and N. Kayagaki. 2021. Dying cells fan the flames of inflammation. Science 374:1076–1080.

Nguyen-Chi, M., B. Laplace-Builhe, J. Travnickova, P. Luz-Crawford, G. Tejedor, Q.T. Phan, I. Duroux-Richard, J.P. Levraud, K. Kissa, G. Lutfalla, C. Jorgensen, and F. Djouad. 2015. Identification of polarized macrophage subsets in zebrafish. Elife 4:e07288.

Nguyen-Chi, M., Q.T. Phan, C. Gonzalez, J.F. Dubremetz, J.P. Levraud, and G. Lutfalla. 2014. Transient infection of the zebrafish notochord with E. coli induces chronic inflammation. Dis Model Mech 7:871–882.

Palecanda, A., and L. Kobzik. 2001. Receptors for unopsonized particles: the role of alveolar macrophage scavenger receptors. Curr Mol Med 1:589–595.

Peeters, P.M., I.M. Eurlings, T.N. Perkins, E.F. Wouters, R.P. Schins, P.J. Borm, W. Drommer, N.L. Reynaert, and C. Albrecht. 2014. Silica-induced NLRP3 inflammasome activation in vitro and in rat lungs. Part Fibre Toxicol 11:58.

Peeters, P.M., T.N. Perkins, E.F. Wouters, B.T. Mossman, and N.L. Reynaert. 2013. Silica induces NLRP3 inflammasome activation in human lung epithelial cells. Part Fibre Toxicol 10:3.

Pfaffl, M.W. 2001. A new mathematical model for relative quantification in real-time RT-PCR. Nucleic Acids Res 29:e45.

Pollard, K.M. 2016. Silica, Silicosis, and Autoimmunity. Front Immunol 7:97.

Rees, D., and J. Murray. 2007. Silica, silicosis and tuberculosis. Int J Tuberc Lung Dis 11:474–484.

Renshaw, S.A., C.A. Loynes, D.M. Trushell, S. Elworthy, P.W. Ingham, and M.K. Whyte. 2006. A transgenic zebrafish model of neutrophilic inflammation. Blood 108:3976–3978.

Riteau, N., L. Baron, B. Villeret, N. Guillou, F. Savigny, B. Ryffel, F. Rassendren, M. Le Bert, A. Gombault, and I. Couillin. 2012. ATP release and purinergic signaling: a common pathway for particle-mediated inflammasome activation. Cell Death Dis 3:e403.

Rushton, L. 2007. Chronic obstructive pulmonary disease and occupational exposure to silica. Rev Environ Health 22:255–272.

Russo, R.C., C.C. Garcia, M.M. Teixeira, and F.A. Amaral. 2014. The CXCL8/IL-8 chemokine family and its receptors in inflammatory diseases. Expert Rev Clin Immunol 10:593–619.

Sayan, M., and B.T. Mossman. 2016. The NLRP3 inflammasome in pathogenic particle and fibre-associated lung inflammation and diseases. Part Fibre Toxicol 13:51.

Schindelin, J., I. Arganda-Carreras, E. Frise, V. Kaynig, M. Longair, T. Pietzsch, S. Preibisch, C. Rueden, S. Saalfeld, B. Schmid, J.Y. Tinevez, D.J. White, V. Hartenstein, K. Eliceiri, P. Tomancak, and A. Cardona. 2012. Fiji: an open-source platform for biological-image analysis. Nat Methods 9:676–682.

Song, L., D. Weng, W. Dai, W. Tang, S. Chen, C. Li, Y. Chen, F. Liu, and J. Chen. 2014. Th17 can regulate silica-induced lung inflammation through an IL-1beta-dependent mechanism. J Cell Mol Med 18:1773–1784.

Spence, R., G. Gerlach, C. Lawrence, and C. Smith. 2008. The behaviour and ecology of the zebrafish, Danio rerio. Biol Rev Camb Philos Soc 83:13–34.

Sztal, T.E., A.A. Ruparelia, C. Williams, and R.J. Bryson-Richardson. 2016. Using Touch-evoked Response and Locomotion Assays to Assess Muscle Performance and Function in Zebrafish. J Vis Exp

Thibodeau, M.S., C. Giardina, D.A. Knecht, J. Helble, and A.K. Hubbard. 2004. Silica-induced apoptosis in mouse alveolar macrophages is initiated by lysosomal enzyme activity. Toxicol Sci 80:34–48.

Torraca, V., C. Cui, R. Boland, J.P. Bebelman, A.M. van der Sar, M.J. Smit, M. Siderius, H.P. Spaink, and A.H. Meijer. 2015. The CXCR3-CXCL11 signaling axis mediates macrophage recruitment and dissemination of mycobacterial infection. Dis Model Mech 8:253–269.

Tyrkalska, S.D., S. Candel, D. Angosto, V. Gómez-Abellán, F. Martín-Sánchez, D. García-Moreno, R. Zapata-Pérez, Á. Sánchez-Ferrer, M.P. Sepulcre, P. Pelegrín, and V. Mulero. 2016. Neutrophils mediate Salmonella Typhimurium clearance through the GBP4 inflammasome-dependent production of prostaglandins. Nat Commun 7:12077.

Tyrkalska, S.D., A.B. Perez-Oliva, L. Rodriguez-Ruiz, F.J. Martinez-Morcillo, F. Alcaraz-Perez, F.J. Martinez-Navarro, C. Lachaud, N. Ahmed, T. Schroeder, I. Pardo-Sanchez, S. Candel, A. Lopez-Munoz, A. Choudhuri, M.P. Rossmann, L.I. Zon, M.L. Cayuela, D. Garcia-Moreno, and V. Mulero. 2019. Inflammasome Regulates Hematopoiesis through Cleavage of the Master Erythroid Transcription Factor GATA1. Immunity 51:50–63 e55.

van der Helm, D., A. Groenewoud, E.S.M. de Jonge-Muller, M.C. Barnhoorn, M.J.A. Schoonderwoerd, M.J. Coenraad, L.J.A.C. Hawinkels, B.E. Snaar-Jagalska, B. van Hoek, and H.W. Verspaget. 2018. Mesenchymal stromal cells prevent progression of liver fibrosis in a novel zebrafish embryo model. Sci Rep 8:16005.

van der Vaart, M., H.P. Spaink, and A.H. Meijer. 2012. Pathogen recognition and activation of the innate immune response in zebrafish. Adv Hematol 2012:159807.

van der Vaart, M., J.J. van Soest, H.P. Spaink, and A.H. Meijer. 2013. Functional analysis of a zebrafish myd88 mutant identifies key transcriptional components of the innate immune system. Dis Model Mech 6:841–854.

Walton, E.M., M.R. Cronan, R.W. Beerman, and D.M. Tobin. 2015. The Macrophage-Specific Promoter mfap4 Allows Live, Long-Term Analysis of Macrophage Behavior during Mycobacterial Infection in Zebrafish. PLoS One 10:e0138949.

Westerfield, M. 2000. The Zebrafish Book. A Guide for the Laboratory Use of Zebrafish Danio* (Brachydanio) rerio. . University of Oregon Press., Eugene, OR.

White, R.M., A. Sessa, C. Burke, T. Bowman, J. LeBlanc, C. Ceol, C. Bourque, M. Dovey, W. Goessling, C.E. Burns, and L.I. Zon. 2008. Transparent adult zebrafish as a tool for in vivo transplantation analysis. Cell Stem Cell 2:183–189.

